# The Lamin A/C Ig-fold undergoes cell density-dependent changes that alter epitope binding

**DOI:** 10.1101/2022.11.22.517482

**Authors:** Melanie Wallace, Gregory R. Fedorchak, Richa Agrawal, Rachel M. Gilbert, Jineet Patel, Sangwoo Park, Matthew Paszek, Jan Lammerding

**Affiliations:** Meinig School of Biomedical Engineering, Cornell University, Ithaca, NY 14850; Weill Institute for Cell and Molecular Biology, Ithaca, NY 14850; Graduate Field of Biophysics, Cornell University, Ithaca, NY 14850

## Abstract

Lamins A/C are nuclear intermediate filament proteins that are involved in diverse cellular mechanical and biochemical functions. Here, we report that recognition of Lamins A/C by a commonly used antibody (JOL-2) that binds the Lamin A/C Ig-fold and other antibodies targeting similar epitopes is highly dependent on cell density, even though Lamin A/C protein levels do not change with cell density. The density-dependent Lamin A/C labeling was distinct from previously reported differential apico-basal labeling, which was independent of cell density. Comparison of the density-dependent labeling effects of antibodies recognizing different Lamin A/C epitopes suggests that the effect is caused by partial unfolding or masking of the C’E and/or EF loops of the Ig-fold in response to cell spreading. Seeding cells on micropatterned surfaces with different areas confirmed that increased cell spreading resulted in reduced Lamin A/C labeling with the JOL-2 antibody. Surprisingly, JOL-2 antibody labeling was insensitive to depolymerization of cytoskeletal filaments or disruption of the Linker of Nucleoskeleton and Cytoskeleton (LINC) complex. Although the density-dependent changes of the Ig-fold did not alter nuclear stiffness or nucleo-cytoskeletal force transmission, they may nonetheless modulate interaction with lamin binding partners and thereby affect cellular functions. Taken together, our results point to a previously unrecognized change in the Lamin A/C Ig-fold that affects recognition by the JOL-2 antibody. These findings are not only important for the interpretation of immunofluorescence data for Lamin A/C, but also raise the intriguing prospect that the conformational changes may play a role in Lamin A/C mediated cellular function.

## Introduction

One way that cells respond to their mechanical environment is through the coupling of the extracellular matrix and cytoskeleton to the nucleus and the corresponding nuclear response to mechanical stress (Wang et al., 2009). The mechanical properties of the nucleus are governed by nuclear lamins and chromatin (Stephens et al., 2017), which respond to matrix elasticity and external forces by modulating the expression and conformation of nuclear lamin proteins (Pajerowski et al., 2007; Swift et al., 2013; Buxboim et al., 2014; Ihalainen et al., 2015; Iyer et al., 2021) and changes in chromatin organization (Furusawa et al., 2015; Stephens et al., 2018b; Nava et al., 2020). Lamin A/C expression scales with tissue stiffness and force application (Swift et al., 2013; Buxboim et al., 2014; Iyer et al., 2021), but despite evidence that the nuclear lamina can stretch in response to mechanical force (Rowat et al., 2006; Pajerowski et al., 2007; Swift et al., 2013), we still have relatively little understanding of how the microenvironment affects lamin conformation and interactions.

Lamins A and C result from alternative splicing of the *LMNA* gene; the two lamin isoforms share the first 566 amino acids, including the immunoglobulin-like (Ig)-fold near the C-terminus. The Lamin A/C Ig-fold serves as the binding sites of countless Lamin A/C binding partners (Donnaloja et al., 2020) and mutations in the Ig-fold can alter Lamin A/C assembly and organization (Shumaker et al., 2005; Verstraeten et al., 2009; Bertrand et al., 2020; Chandrayee Mukherjee et al., 2022). The Lamin A/C Ig-fold can respond to mechanical stimuli through conformational changes that allow for the exposure of a cryptic cysteine reside, Cys^522^, in response to shear stress (Swift et al., 2013) and the selective binding of antibodies in response to actin-mediated cell spreading or force application to the nucleus (Ihalainen et al., 2015). Many Lamin A/C Ig-fold binding partners that are important for nucleo-cytoskeletal coupling, chromatin organization, and regulation of gene expression interact with the particular Ig-fold region subject to mechanically induced conformational changes (Swift et al., 2013; Ihalainen et al., 2015), including actin, DNA, emerin, LAP2α, and SUN1/2 (Donnaloja et al., 2020). Perhaps even more intriguingly, the Lamin A/C Ig-fold is a hotspot for *LMNA* mutations that cause several diseases collectively referred to as “laminopathies” (Scharner et al., 2014). These *LMNA* mutations may cause structural change or destabilize the Ig-fold (Shumaker et al., 2005; Qin et al., 2011; Dutta et al., 2018), which could affect Lamin A/C function and interaction with binding partners. Therefore, it is imperative to elucidate the nature of Lamin A/C Ig-fold conformational changes and/or binding partner interactions in response to microenvironmental changes.

Here, we report that the Lamin A/C Ig-fold undergoes conformational changes or epitope masking in response to cell seeding density, as evidenced by the cell density-dependent binding of a commonly used anti-Lamin A/C antibody that binds the Ig-fold (JOL-2 antibody, referred to here as LAC-Ig1). Importantly, Lamin A/C levels did not change with cell density, as confirmed by immunoblotting, Lamin A/C antibodies targeting different epitopes, and cells with fluorescently labeled endogenous Lamin A/C. Based on available epitope mapping of the anti-Lamin A/C antibodies used in this study, we propose two non-mutually exclusive hypotheses to explain these findings: (1) the LAC-Ig1 antibody binds the C’E loop of the Ig-fold, which partially unfolds in response to tension on the nucleus, in line with previous results showing unfolding of the C’E loop in response to mechanical stress (Swift et al., 2013), and/or (2) cell spreading alters interactions of the Lamin A/C Ig-fold with its binding partners, for example, by changes in the secondary structure, post-translational modifications, or its accessibility within the lamin filament network, resulting in masking of the epitope recognized by the LAC-Ig1 antibody. Unlike in previous studies, the specific change in antibody recognition reported here was independent of cytoskeletal forces acting on the nucleus, as neither actin, microtubules, nor LINC complex disruption abolished the cell density dependent JOL-2 labeling. Although the precise mechanisms and functional consequences of the change in epitope recognition remains to be elucidated, these findings add to a growing body of evidence (Swift et al., 2013; Ihalainen et al., 2015) that the Lamin A/C Ig-fold is subject to conformational changes or other regulation in vitro, which may govern interaction of Lamin A/C with specific binding partners and thus modulate cellular function.

## Materials and Methods

### Cell culture

Immortalized human fibroblast, HeLa, and MDA-MB-231 cells were cultured in tissue culture-treated flasks in Dulbecco’s Modified Eagle Medium (DMEM; Gibco) plus 10% Fetal Bovine Serum (FBS) and 1% Penicillin-Streptomycin (P/S), referred to here as D-10 medium. For imaging and microharpoon experiments, cells were dissociated as single cells using 0.25% Trypsin and resuspended in D-10 medium. Cells were then counted and seeded in a serial dilution, according to Table 1 for imaging experiments, on sterile, fibronectin-coated (1 µg/mL) coverslips to achieve a range from low seeding densities, in which there are rare cell-cell contacts, to very confluent seeding densities. Cells were allowed to adhere to coverslips or plates for 24 hours prior to experimentation, and were visually inspected to ensure a uniform seeding density was achieved. Areas in which the cell density substantially deviated from the expected cell density were excluded from the image acquisition. Alternatively, cells were seeded at the same density and collected at different time points to obtain the desired density.

**Table 1.** Cell seeding density for experiments using human fibroblast and HeLa cells. Cell seeding densities ranged from fully confluent with numerous cell-cell contacts to low seeding densities with extremely low confluency and rare cell-cell contacts. Cell numbers refer to the number of cells per well in a 6-well plate atthe time of seeding.

| Dilution | # of Cells | Cell Density (#/cm<sup>2</sup>) | % Confluency |
| --- | --- | --- | --- |
| 1.5× | 60,000 | 31,579 | 100% |
| 1.0× | 40,000 | 21,053 | 80% |
| 0.5× | 20,000 | 10,526 | 40% |
| 0.25× | 10,000 | 5,263 | 20% |
| 0.125× | 5,000 | 2,632 | 10% |
| 0.0625× | 2,500 | 1,316 | 5% |

### CRISPR knock-in generation

The endogenous *LMNA* gene was targeted using lentiCRISPR v2 directed at exon 1 site 2 (spacer sequence: CATCGACCGTGTGCGCTCGC). Gibson assembly was used to generate the repair plasmid within a DTA killer gene backbone (PGKdta bpa) and consisted of an mNeonGreen fluorophore cloned from a pCDH mNeonGreen Prelamin A plasmid. The use of site directed mutagenesis to induce silent mutations in the repair construct prevented secondary cutting by CRISPR. MDA-MB-231 cells were plated in a 6-well plate, transduced with recombinant adeno-associated virus (rAAV) (Addgene, pAAV TBG FFLuc) supernatant, and allowed to incubate for 48 hours. Cells were split to reduce the density and then transiently transfected with the lentiCRISPR v2 E1E2 plasmid consisting of an insertion with Blasti-P2A-mNeonGreen. Cells were placed under high selection with blasticidin. rAAV-modified MDA-MB-231 cells were sorted for mNeonGreen signal using the Cornell Imaging Facility FACS. Non-rAAV-modified cells were used as controls. After two rounds of sorting, mNeonGreen-positive cells (mNG-LMNA) were enriched from <10% to almost 100%. While some populations were left as heterogeneous, others were sorted based on intensity (high and low expression) to test distinct subpopulations. For clonal selection, cells were seeded at very low density in 10 cm^2^dishes and grown for several weeks. Clonal rings and vacuum grease were used to select individual colonies for expansion. Both heterogenous and clonal populations were generated and used in experiments, as indicated in the figure legends.

### Generation of cells with inducible LINC complex disruption

A tet-on inducible dominant negative nesprin-4 KASH domain construct fused with GFP (GFP-KASH4-TETON) described previously (Roux et al., 2009) was cloned into a pRetroX-TightPur vector. Ectopic expression of KASH4 displaces endogenous nesprin-2G and nesprin-3 from the nuclear envelope (Roux et al., 2009). For viral production, 293 GPG cells were transfected using Lipofectamine and the constructs of interest and viral particles were collected from day 3 to day 7. Human fibroblasts were first stably modified with an rtTA-Advanced transactivator required to express the gene of interest in the response vector, then with the construct with the gene of interest. Polybrene at a concentration of 8 μg/mL was used to enhance the infection. For experiments testing the effect of LINC complex disruption, cells expressing the KASH4 or control constructs were treated with 500 ng/mL Doxycycline for a minimum of 24 hours in order to induce expression of the construct of interest.

### DNA isolation and PCR gel

DNAzol Reagent (Invitrogen) was added to cells seeded in a 6-well plate and genomic DNA was isolated according to the manufacturer’s instructions. The following primer pairs were used to evaluate proper insertion of the mNG tag into the correct location in the genome: mNeonGreen_seq_F = 5’ CCCAACGACAAAACCATCAT 3’, *LMNA* downstream_R = 5’ TGCAGTTGAGTAGGGTGGG3’, LMNA upstream_F = 5’AACTCCTTGATCCCTGGCC3’ and LHA (left homology arm) extract_R = 5’ACCGCCAAGCGATCAT 3’. Amplicons were amplified using Phusion DNA Polymerase (NEB). PCR products were run on an agarose gel using standard lab protocols and imaged.

### Immunoblotting

Cells were either lysed in equal cell numbers in Laemmli sample buffer (Bio-Rad) containing 0.1 M of Dithiothreitol (DTT) after trypsinization or in RIPA buffer supplemented with proteinase inhibitor (Roche). Protein content was measured in the RIPA cell lysates using a standard Bradford assay. All samples were heat-denatured (5 min at 95°C) in Laemmli sample buffer and separated by SDS-PAGE (Invitrogen). Proteins were transferred onto PVDF membranes (Millipore, IPVH00010) using semi-dry transfer method (Bio-Rad). Immuno-detection was carried out with the following primary antibodies: anti-Lamin A/C (Abcam #40567 for cell density studies, Santa Cruz Biotechnologies sc-6215 for mNG-LMNA knock-in, dilution 1:2000), anti-β-tubulin (Sigma, T5168, dilution 1:4000). Blots were probed with HRP-conjugated antibodies (Biorad, dilution 1:1000; Jackson Immuno Research, dilution: 1:10000). See Supplementary Table for further details.

### Activation and inhibition of retinoic acid pathway

Cultured cells were treated with all-trans retinoic acid (RA, 1 µM, Fisher Scientific), high affinity pan-RAR antagonist AGN-193109 (AGN, 1 µM, Santa Cruz Biotechnology), or vehicle control (0.15% EtOH, 0.15% DMSO in media with 10% FBS) for 48 hours.

### Pharmaceutical inhibitor studies

Human fibroblasts were seeded, at high (10,000 cells per well) and low (1,000 cells/well) density, into 96-well plates, allowed to adhere, and treated for 24 hours (unless otherwise noted) prior to fixation. The following treatments were used: a histone deacetylase inhibitor, Trichostatin A (TSA, 200 nM), a microtubule depolymerizing agent, Nocodazole (75 nM, 24h prior to fixation), or an actin depolymerizing agent, Cytochalasin D (CytoD, 10 µM, 60 minutes prior to fixation). A separate human fibroblast line stably modified to express an inducible dominant-negative KASH (DN-KASH) construct was treated with 500 ng/mL Doxycycline for 24 hours prior to fixation to disrupt nucleo-cytoskeletal coupling.

### Immunofluorescence staining

Cells were passaged as previously described onto fibronectin-coated coverslips. The next day, media was either changed to fresh D-10 or drug-treated D-10, and then cells were washed with 1× PBS and fixed in warmed Paraformaldehyde (2% or 4%) for 10 minutes or on ice in cold 1:1 Methanol and Acetone for 15 minutes. Cells were then washed with 1× PBS and blocked in 3% BSA with 0.1% Triton-X 100 (Thermo-Fischer) and 0.1% Tween (Sigma) in PBS for one hour at room temperature. Primary antibodies (Table S1) were prepared in blocking solution and incubated overnight at 4°C. Cells were then washed with a solution of 0.3% BSA with 0.1% Triton-X 100 and 0.1% Tween in PBS and stained with AlexaFluor secondary antibodies (1:250; Invitrogen) for 1 hour at room temperature. DAPI (1:1000, Sigma) was added for 10 minutes at room temperature, and cells were washed with PBS before imaging. See Supplementary Table for details on antibody use and concentrations.

### Micropatterning experiments

Micropatterns were generated based on an existing protocol (Tan et al., 2009). In brief, 4-inch glass wafers (0211#1 1/2, Precision Glass & Optics) were cleaned with 1:1 methanol and hydrochloric acid for 15 minutes, followed by 1 minute in a plasma cleaner (Harrick Plasma, PDC-001). A methyl ether polyethylene glycol with a reactive tri-ethoxy silane (mPEG-Silane MW 5k, Creative PEGWorks) was coated on the wafer, and a thin film of parylene C (diX-C, Uniglobe Kisco) was deposited on the PEG layer via chemical vapor deposition system (PDS 2010 LABCOTER deposition system) to 800 nm thickness. The thickness of parylene was measured with profilometer (Veeco Dektak 6M Profilometer), and the parylene-coated glass wafer was patterned using standard photolithography. A photoresist layer (S1827) was spun on top of the parylene layer and desired patterns were exposed on the photoresist using 405 nm ultraviolet light (SUSS MA6-BA6 Contact Aligner). After the photoresist was developed, the parylene were etched by oxygen plasma (Oxford PlasmaLab 80+). Residual photoresist was removed by rinsing the wafer with n-methyl-2-pyrrolidone (NMP). Each pattern on the wafers was diced into 10 mm by 10 mm chips, consisting of circle features with diameters of either 20 µm or 40 µm. The chips were sterilized with UV for 15 minutes. 0.1 mg/mL of fluorescently labeled human fibronectin in phosphate-buffered saline (PBS) was incubated with the chips for 1 hour at 37°C. The chips were then rinsed twice with PBS to remove unbound fibronectin and the parylene film was mechanically peeled off with sterile tweezers.

Human fibroblasts were then dissociated as single cells using 0.25% Trypsin, resuspended in D-10 medium, and counted. 160,000 cells suspended in 500 μL of media were plated onto each pattern in a 24-well plate and shaken to distribute cells across the surface. Plates were shaken every five minutes to prevent adherence of cells off of the micropatterns. Once cells were firmly attached to the patterns (20 minutes for 40-μm patterns or 30 minutes for 20-μm patterns), the media was removed and patterns were gently washed with 1× PBS. Devices were then transferred to a fresh well to prevent any remaining cells from adhering off of the patterns and incubated at 37°C for 1.5 hours after seeding to allow cells to spread. Devices were subsequently fixed and stained as specified for immunofluorescence experiments.

### Image acquisition and analysis

Flasks and coverslips in cell culture plates were imaged with a Zeiss LD Plan-Neofluar 20× phase contrast objective (NA = 0.4) on an inverted Zeiss Axio Observer Z1 epifluorescence microscope equipped with a Zeiss Colibri 7 solid state LED illumination system and a Hamamatsu ORCA Flash 4.0V2 sCMOS camera. Exposure times were set for each experiment based on the lowest cell seeding density, which had the highest Lamin A/C immunofluorescence intensity, to avoid saturation. All experimental groups were then imaged at the identical settings. To quantify the intranuclear distribution of Lamin A/C labeling, apico-basal distribution of Lamin A/C labeling, and nuclear height, cells on coverslips were imaged as stacks using a 63× Plan-Apochromat oil immersion objective (NA = 1.4) on a Zeiss laser scanning confocal microscope equipped with photomultiplier tubes as detectors (LSM700). Pixel dwell times and amplification gain were set for each experiment based on the lowest cell seeding density to avoid saturation.

Mean fluorescence intensity and area of each nucleus was obtained using custom MATLAB (Mathworks) scripts to threshold nuclei. Similarly, actin fluorescence intensity was computed by dividing the total actin fluorescence intensity in an image by the number of nuclei in the image. For the majority of experiments, fluorescence intensity values were normalized for each experiment to account for variability in immunofluorescence experiments and image acquisition.

Cell area was computed as an average over an image from actin immunofluorescence images. Thresholded actin images were converted to binary images, the total area covered by cells computed, and then divided by the number of nuclei in the image.

For micropattern experiments taken as image stacks, thresholding was performed, and then the total fluorescence intensity of each nucleus was summed to account for differences in nuclear size. The total summed fluorescence intensity of each nucleus was then averaged across all nuclei at a given experimental condition and normalized to the average summed fluorescence intensity for the 20-μm patterns.

Nucleoplasmic and peripheral fluorescence intensities of the LAC-Ig1 signal were computed from high-resolution confocal z-stacks acquired of cells seeded at either high (1.0×) or low (0.0625×) cell density and immunofluorescently labeled with the LAC-Ig1 antibody. For each nucleus, the z-position for which the x-y plane dissects the center of the nucleus was identified, and a line was drawn across the major axis of the nucleus to obtain the Lamin A/C fluorescence intensity values along with position along the line. Fluorescence intensity profiles were trimmed such that the first and last values in the profile corresponded to the outer edges of the NE. Positions across each nucleus were then normalized to a scale of 0 to 1 to account for differences in nuclear size. Fluorescence intensity values and the corresponding normalized positions for each nucleus for each seeding density were then read into a custom R script for analysis (script available upon request). Fluorescence intensity profiles across all nuclei in a cell line were averaged and normalized to the area under the curve to account for variations in fluorescence intensities due to cell seeding density or staining conditions. The boundary of the nuclear lamina was determined by the inflection point of the average fluorescence intensity profile of all cell lines with the nuclear lamina defined as the peripheral regions with normalized nuclear positions of <0.16 and >0.84, and the nucleoplasmic region for positions between normalized positions of 0.16 and 0.84. For each nucleus, the fluorescence intensity of nucleoplasmic lamins was then determined by taking the 95^th^ percentile of fluorescence intensity values in the nuclear lamina region (to avoid influence of spurious pixel intensity values), and the fluorescence intensity of nucleoplasmic lamins was determined by taking the average value of fluorescence intensities within the nucleoplasm region. The ratio of nucleoplasmic to peripheral lamins was then computed for each nucleus and plotted.

To quantify the apico-basal distribution of Lamin A/C and nuclear height, cells on coverslips were imaged as stacks with a 63× Plan-Apochromat oil immersion objective (NA = 1.4) on a Zeiss laser scanning confocal microscope equipped with photomultiplier tubes as detectors (LSM700). Exposure time was set for each experiment based on the lowest cell seeding density to avoid saturation. Airy units were set to 1.0 for all images. Apico-basal polarization analysis was completed with a custom MATLAB (Mathworks) script in which the image stacks of cells labeled for Lamin A/C were thresholded, and the fluorescent intensity in each image slice was computed at the centroid of each nucleus. A measure of apico-basal polarization was computed by taking the average centroid intensity of the bottom half of image slices over the average centroid intensity of top half of image slices.

### Biophysical assays

For micropipette aspiration experiments, a confluent T75 flask was split 1:12 (∼87,500 cells) into a T25, T75 and T150 flask 48 hours prior to experimentation to establish three different densities (low, medium, high). Micropipette aspiration was performed on human fibroblasts according to a previously described protocol (Davidson et al., 2019). A pressure differential of 1.0 psi and 0.2 psi at the inlet and outlet reservoirs drove the flow of cells through the device. Images were acquired every 5 seconds using an inverted microscope for a minimum of 60 seconds. Nuclear protrusion length was calculated using a custom MATLAB script available upon request.

For microharpoon studies, human fibroblasts were seeded 24 hours prior to experimentation. The microharpoon experiment was performed as described previously (Fedorchak and Lammerding, 2016). The microharpoon was inserted ≈5 µm from the edge of the nucleus and pulled 16 μm at 4 μm/s. Images were acquired with a 40× Zeiss EC Plan-Neofluar objective (NA = 0.75, Ph2) every 5 seconds. Average nuclear strain and centroid displacement were calculated using a custom MATLAB script available upon request.

### Lamin A/C Ig-fold structure

Lamin A/C Ig-fold 3D structures were rendered from the 1IVT Ig-fold molecular structure (Krimm et al., 2002) using the VMD Molecular Graphics Viewer (http://www.ks.uiuc.edu/Research/vmd/; Humphrey et al., 1996)

### Statistics

All results were acquired from two to seven independent experiments. To account for inherent cell-to-cell variability or variation in local seeding density, and to remove any unintentional bias, we did not remove any outliers. Thus, violin plots presented in this manuscript contain all experimental data points of single cell/nuclei measurements. Numeric datasets in which whole images or single cells or nuclei were analyzed were tested for normality, and data not following a normal distribution were linearized either by taking the natural log or the square root, whichever achieved better normalization. Either a Student’s *t*-test (for two groups) or one-way analysis of variance (ANOVA) with multiple comparisons (for more than two groups) was then performed in GraphPad Prism, using Tukey’s correction for multiple comparisons.

## Results

### Lamin A/C epitope immunolabeling changes in a density-dependent manner

The Lamin A/C Ig-fold is known to undergo mechanosensitive conformational changes in response to mechanical stimuli (Swift et al., 2013; Ihalainen et al., 2015), which may be indicated by differential binding of Lamin A/C antibodies recognizing epitopes within the Ig-fold (Ihalainen et al., 2015). Furthermore, mechanosensitive changes in the Lamin A/C organization could alter binding of Lamin A/C with other proteins, which could result in masking of specific epitopes. We hypothesized that variations in cell seeding density that correspond to changes in cell spreading and cell-cell contacts could alter nuclear tension and result in mechanosensitive conformational changes and antibody recognition. We seeded human fibroblasts in a serial dilution such that the lowest cell seeding density (0.0625×, resulting in ≈1,315 cells/cm^2^) corresponded to low confluency, large cell spread area, and few cell-cell contacts, and the highest seeding density (1.5×, resulting in ≈31,580 cells/cm^2^) corresponded to nearly 100% confluency, low cell spread area, and many cell-cell contacts (Fig. 1A, C). Cell densities were visually inspected to ensure uniform cellular distribution. We performed immunofluorescence labeling for Lamin A/C across six cell seeding densities in immortalized human fibroblasts using the JOL-2 antibody (LAC-Ig1), and quantified Lamin A/C fluorescence intensity in single nuclei to account for any potential variation due to small local variations. We observed a striking difference in LAC-Ig1 fluorescence intensity across cell seeding densities (Fig. 1B). LAC-Ig1 fluorescence intensity per nucleus varied inversely with cell seeding density, and cells seeded at lower densities had significantly increased LAC-Ig1 fluorescence intensity compared to cells seeded at higher densities (Fig. 1D, data normalized to the values at the highest seeding density, 1.5×). The two lowest seeding densities, 0.0625× and 0.125×, exhibited similarly increased LAC-Ig1 fluorescence intensity, which could indicate that at the lowest seeding densities, cells are already very spread apart and have little cell-cell contacts, thus resulting in more similar cell spreading areas and/or cytoskeletal conformations.

**Figure 1.**
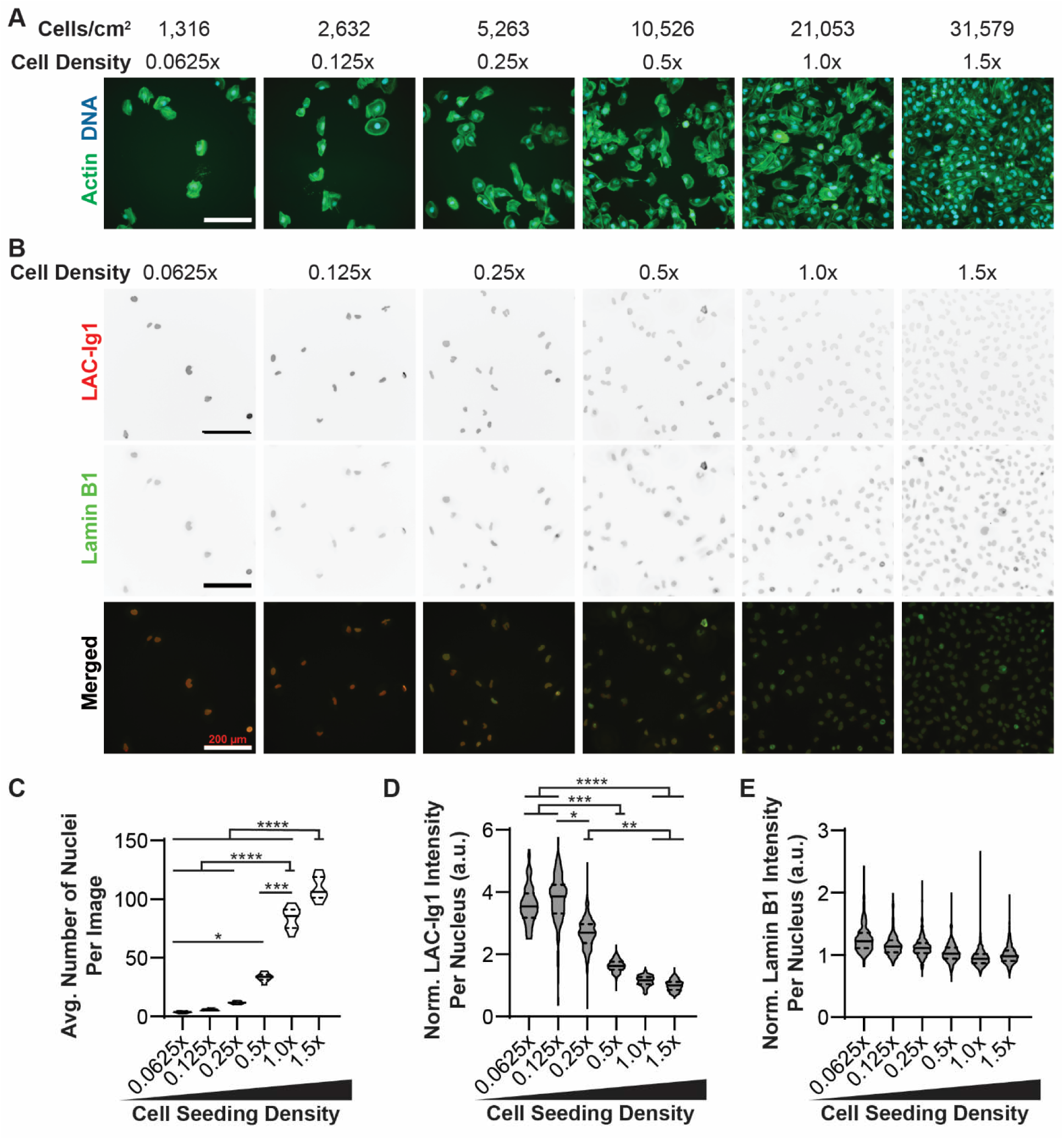
Lamin A/C epitope expression changes in a density-dependent manner in human fibroblasts. (**A**) Representative images of human fibroblasts labeled for actin seeded in a serial dilution showing that at low cell seeding densities, cells are at a low confluency with only few cell-cell contacts, and at the highest cell seeding densities, cells are in a nearly confluent monolayer with numerous cell-cell contacts. (**B**) Representative images of human fibroblasts seeded roughly in a serial dilution and immunofluorescently labeled for Lamin A/C (red) with the JOL-2 antibody (LAC-Ig1) or for Lamin B1 (green). Scale bars: 200 μm. (**C**) Quantification of the average number of human fibroblast nuclei per image section as a function of cell seeding density. Data based on *N* = 30 images per group from three independent experiments. (**D, E**) Corresponding quantification of Lamin A/C (D) and Lamin B1 (E) immunofluorescence intensity of human fibroblasts as a function of cell seeding density, normalized to the highest cell seeding density, 1.5×. *, *p* < 0.05; **, *p* < 0.01; ***, *p* < 0.001; ****, *p* < 0.0001. *N* > 100 nuclei per group for fluorescence intensity, across three independent experiments.

To confirm that density-dependent immunolabeling with the LAC-Ig1 antibody was not the result of a specific fixation method, we performed the experiments with two alternative fixation methods, 2% paraformaldehyde (PFA) or 1:1 Methanol:Acetone. We observed the same inverse correlation of LAC-Ig1 immunolabeling and cell density with both methods (**Fig. S1**). To rule out that the labeling effect was due to general cell density-dependent differences in antibody depletion or accessibility, we quantified Lamin B1 fluorescence intensity across the same cell seeding densities. Despite its intracellular localization similar to Lamin A/C, Lamin B1 immunolabeling did not show any density-dependent changes in fluorescence intensity (Fig. 1B, E), indicating that the cell density effect was specific to Lamin A/C and labeling with the LAC- Ig1 antibody.

To test whether density-dependent immunolabeling with Lamin A/C-Ig-1 was conserved in other human cell types, we repeated the experiment with HeLa cells, an immortalized epithelial adenocarcinoma cell line, and with MDA-MB-231 cells, a triple-negative epithelial adenocarcinoma cell line. We observed a similar inverse relationship between LAC-Ig1 immunofluorescence intensity and cell seeding density in HeLa cells (**Fig. S2A, B**), whereas immunofluorescence intensity for Lamin B1 was independent of cell density (**Fig. S2C**). Similar findings were obtained in the MDA-MB-231 cell lines (**Fig. S3**). For these cells, we explored an additional variation: instead of seeding cells simultaneously at different cell densities, cells were seeded at a similar low density, and then fixed and immunofluorescently labeled at different time points, after having reached cell densities comparable to the other experiments. This approach ensured that all cells were fixed while actively proliferating, thereby reducing potentially confounding effects from cell-contact inhibition. As in the other experiments, immunolabeling of Lamin A/C with the LAC-Ig1 antibody produced the same inverse relationship between LAC- Ig1 fluorescence intensity and cell seeding density, with a significant increase in LAC-Ig labeling at low seeding densities compared to high seeding densities (**Fig. S3B**). Collectively, these results indicate that LAC-Ig1 density-dependent immunolabeling is consistently observed across multiple human cell lines. Since the LAC-Ig1 antibody specifically detects human Lamin A/C, we did not investigate cells from other species. Instead, we focused primarily on human fibroblasts as a representative cell line for future experiments.

### LAC-Ig1 density-dependent variation is not due to changes in protein expression

To test whether density-dependent Lamin A/C immunofluorescence labeling could be due to changes in protein expression levels, we performed immunoblot analysis for Lamin A/C of cells seeded at different densities. We found no significant differences in Lamin A/C protein levels (Fig. 2A), indicating that density-dependent LAC-Ig1 variation was not caused by changes in Lamin A/C protein levels.

**Figure 2.**
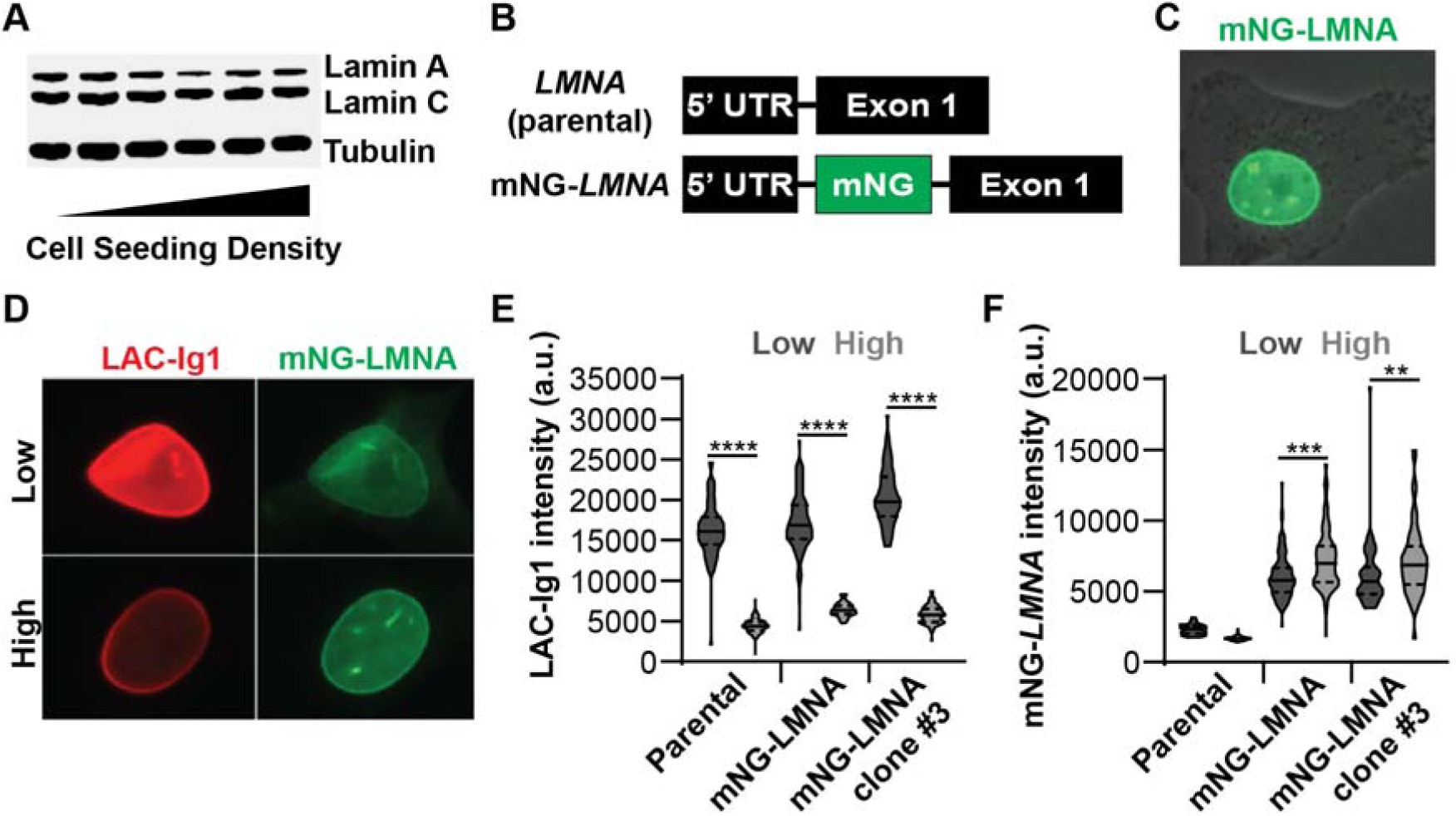
Density-dependent Lamin A/C immunolabeling is not due to altered protein levels. (**A**) A Western Blot shows that human fibroblasts Lamin A/C protein levels are similar at different cell seeding densities. (**B**) An mNeonGreen (mNG) fluorescent tag was inserted into the *LMNA* gene (mNG-*LMNA*) in between the 5’ untranslated region (UTR) and exon 1. (**C**) mNG-Lamin A/C was highly expressed and localized only to the nucleus in MDA-MB-231 cells, as expected. (**D**) Representative images of MDA-MB-231 cells expressing mNG-LMNA (green) and immunofluorescently labeled for Lamin A/C using the LAC-Ig1 antibody (red). (**E**) LAC-Ig1 immunofluorescence intensity was significantly reduced at high cell densities in parental, heterogeneous mNG-LMNA, and a clonal mNG-LMNA cell line. ****, *p* < 0.0001. *N* > 60 nuclei per group from two independent experiments. (**F**) mNG-LMNA fluorescence signal intensity did not follow the same density-dependent effect as the LAC-Ig1 immunofluorescence, exhibiting only some small, significant *increase* in signal intensity at *high* cell densities. **, *p* < 0.01; ***, *p* < 0.001. *N* > 60 nuclei per group from two independent experiments.

To confirm these results using an independent method, we generated MDA-MB-231 cells in which one of the endogenous *LMNA* alleles was fluorescently tagged (Fig. 2B). We used CRISPR/Cas9 to insert an mNeonGreen (mNG) sequence into the genomic locus corresponding to the N-terminus of endogenous Lamin A/C, thus producing N-terminally tagged Lamin A/C (mNG-LMNA) (**Fig. S4A**) and enabling us to visualize directly changes in Lamin A/C protein expression. We validated correct insertion by PCR (**Fig. S4B**), immunoblotting (**Fig. S4C**), and immunofluorescence (Fig. 2C), demonstrating that mNG-LMNA was properly expressed and localized to the nucleus (Fig. 2C). Notably, the expression levels of the tagged-Lamin A/C were low compared to non-tagged Lamin A/C (**Fig. S4C**), suggesting that only a single allele out of the three *LMNA* alleles in the MDA-MB-231 cells was tagged. In addition, we confirmed that the mNG-LMNA modification did not alter nuclear stiffness (**Fig. S4D**).

We then seeded heterogeneous mNG-LMNA cells, a clonal mNG-LMNA cell line, and the parental, non-modified controlled cells at either high or low cell seeding densities and quantified the fluorescence intensity of immunolabeling with the LAC-Ig1 antibody (Fig. 2D, E) and the mNeonGreen signal from the mNG-LMNA construct (Fig. 2D, F) at each density. Consistent with our earlier findings, the LAC-Ig1 fluorescence intensity was significantly increased in cells at low seeding densities compared to those in high seeding densities (Fig. 2E). In contrast, we did not observe any increase in the mNeonGreen fluorescence signal in the mNG- LMNA cells, confirming that the increase in the LAC-Ig1 signal was not due to increased Lamin A/C levels. In fact, we observed a small but significant *increase* in mNG-LMNA expression at *high* seeding densities for mNG-LMNA expressing cells (Fig. 2F), although the scale of these changes was much smaller compared to those observed with the immunolabeling with the LAC- Ig1 antibody. Collectively, these results confirm that the density-dependent change in LAC-Ig1 immunolabeling was not due to changes in Lamin A/C protein expression. Rather, density-dependent immunolabeling likely results from conformational changes of the Lamin A/C Ig-fold that alter the accessibility or recognition of certain protein domains and causes changes in epitope binding.

### Lamin A/C density-dependent epitope binding does not correspond to changes in nuclear stiffness

Since Lamin A/C is a key determinant of nuclear mechanical properties (Lammerding et al., 2006; Stephens et al., 2018a), we asked whether density-dependent LAC-Ig1 epitope binding, which may be indicative of conformational changes, altered organization, or binding partner interactions in the Lamin A/C protein or network, is reflective of changes in nuclear mechanics. To measure nuclear mechanics, we quantified nuclear stiffness of cells that had been seeded at different cell densities using a high-throughput micropipette aspiration assay (Davidson et al., 2019). In this assay, cells were detached immediately prior to the experiments and perfused into the microfluidic device. A pressure gradient was then applied across a series of microfluidic channels, the nucleus was aspirated into a narrow channel, and the length of the nuclear protrusion into the channel was used as a proxy for nuclear stiffness (Fig. 3A). Notably, we did not observe any density-dependent changes in nuclear protrusion length into the channel (Fig. 3B), indicating that nuclear stiffness did not change with cell seeding density.

**Figure 3.**
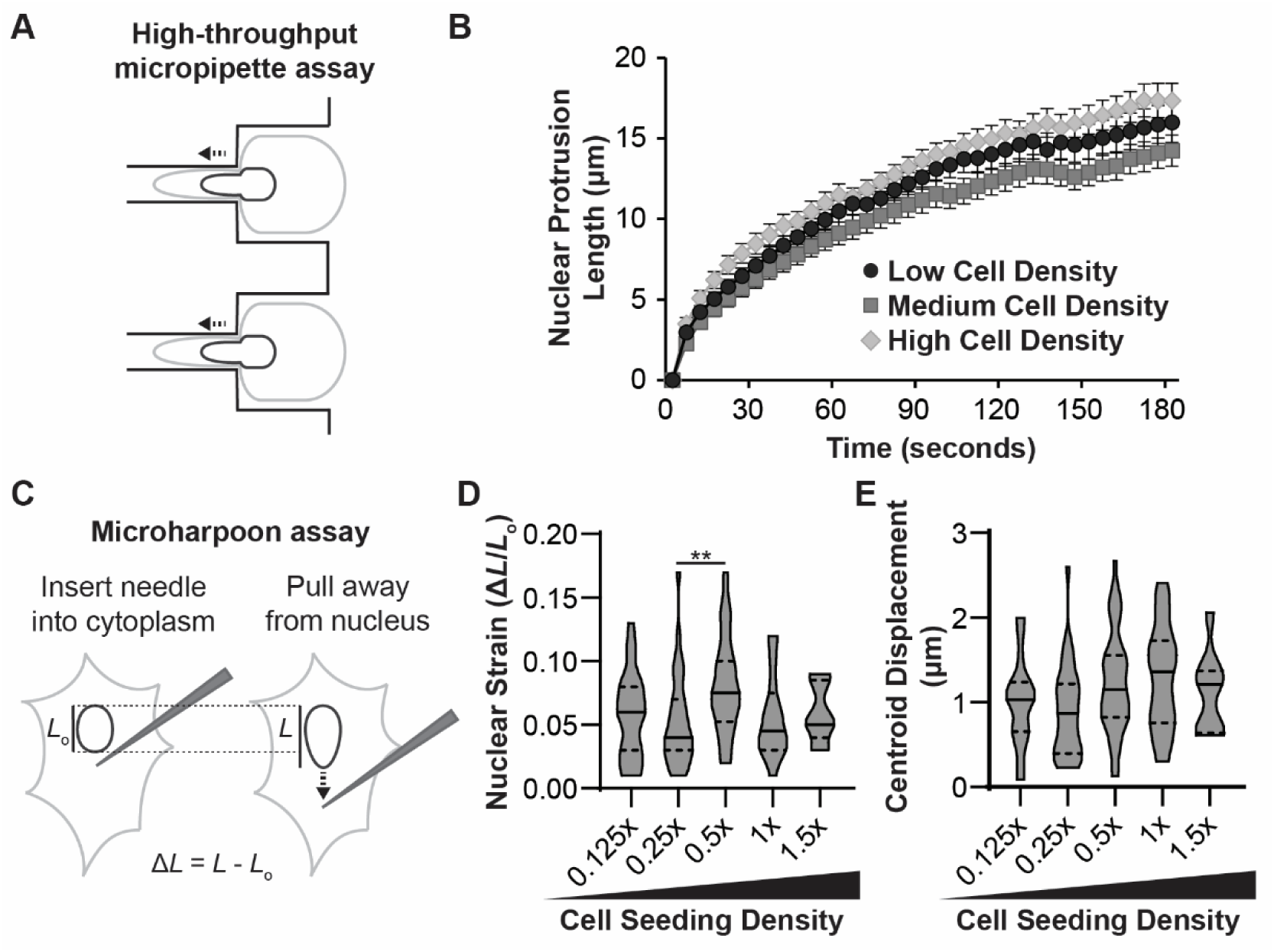
Density-dependent Ig-fold conformational changes do not correspond to changes in nuclear mechanical properties. (**A**) Schematic depiction of cell and nuclear deformation in the microfluidic micropipette aspiration assay. The nuclear protrusion length into the small channel is used as an indicator of nuclear stiffness. (**B**) Cells seeded at low, medium, and high densities had similar nuclear protrusion lengths, indicating that nuclear stiffness did not vary with cell seeding density. (**C**) Schematic depiction of the microharpoon assay, in which a microneedle is inserted into the cytoplasm near the nucleus and pulled away for a fixed distance. Nuclear strain and displacement are used to infer nuclear stiffness and nucleo-cytoskeletal coupling, respectively. Corresponding quantification of (**D**) nuclear strain and (**E**) nuclear centroid displacement for cells seeded at different cell densities revealed no significant trends across cell seeding densities. **, *p* < 0.01. *N* > 50 nuclei per group for micropipette, across three independent experiments. *N* ≥ 9 nuclei per group for microharpoon, from three independent experiments.

To address the possibility that changes in lamin conformation, organization, or binding partner interaction do not persist following cell detachment and suspension, we used an alternative approach to measure nuclear mechanics in adherent cells seeded at different cell densities. In this ‘microharpoon assay’, a microneedle is inserted into the perinuclear cytoskeleton and pulled away from the nucleus with precisely controlled speed (Fig. 3C); information about nuclear stiffness and nucleo-cytoskeletal coupling can then be inferred from the nuclear deformation (nuclear strain) and movement (centroid displacement), respectively

(Fedorchak and Lammerding, 2016). While we observed some small variations in nuclear strain in cells seeded at intermediate cell seeding densities (Fig. 3D), neither nuclear strain nor nuclear centroid displacement (Fig. 3E) showed any correlation with cell seeding density. Together, these results indicate that density-dependent changes in the Lamin A/C network that alter LAC- Ig1 binding do not correspond to significant changes in the mechanical properties of the nucleus.

### Lamin A/C density-dependent epitope binding corresponds to cellular and nuclear area

Both cell spread area and cell-cell contacts vary with cell seeding density and could play a role in the mechanical conformation of the cell. Cell spread area in particular has been shown to govern Lamin A/C Ig-fold conformational changes, with increased cell spreading resulting in a differential apico-basal conformation of the nuclear lamina that allows some Lamin A/C Ig-fold targeting antibodies to preferentially bind on the apical surface of the nucleus (Ihalainen et al., 2015). To assess to what extent cell spread area varies with the cell seeding densities presented here, we quantified the spread area of human fibroblasts seeded in a serial dilution. Human fibroblasts seeded at the lowest three cell seeding densities (0.0625×, 0.125×, 0.25×) had significantly increased cell spread areas compared to cells seeded at high densities (Fig. 4A). We also quantified nuclear cross-sectional area in the human fibroblasts seeded in a serial dilution, since increased nuclear area in highly spread cells could indicate increased tension or compression of the nuclear lamina. Similar to cell cross-sectional area, human fibroblasts seeded at the low cell densities had significantly increased nuclear area compared to cells seeded at high seeding densities (Fig. 4B). Together, data suggest that density-dependent LAC-Ig1 immunolabeling may correlate to both cellular and nuclear spread area, which is in line with previous studies (Ihalainen et al., 2015). However, whereas human fibroblasts showed an increase in nuclear cross-sectional area with decreasing cell density (Fig. 4B), this trend was not or only partially observed in HeLa cells (**Fig. S5A**) and MDA-MB-231 cells (**Fig. S5B**), even though these cells show a similar cell density dependent LAC-Ig1 immunolabeling (**Figs. S2, S3**). These data suggest that cell spreading, and not nuclear flattening determine the observed LAC-Ig1 labeling effect.

**Figure 4.**
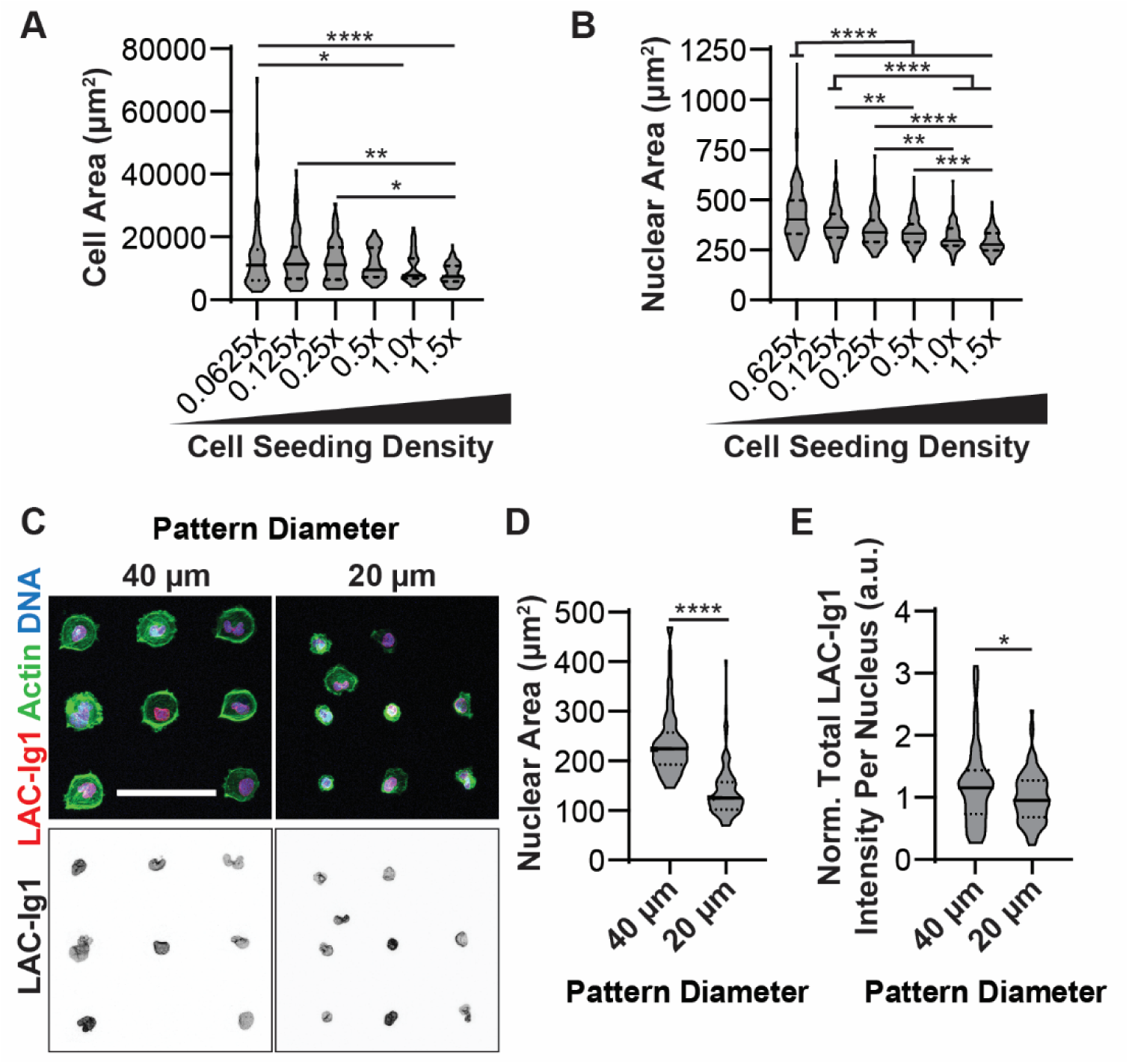
Changes in cell spread area are sufficient to cause differential binding of LAC-Ig1 antibody. Human fibroblasts seeded at low cell seeding densities exhibited (**A**) increased cell spread area and (**B**) increased nuclear cross-sectional area compared to high cell seeding densities. (**C**) Representative immunofluorescence images of human fibroblasts seeded on either 20- or 40-μm diameter micropatterns. Scale bar: 100 µm. Human fibroblasts seeded on 40-μm diameter micropatterns had significantly increased (**D**) nuclear area and (**E**) total LAC-Ig1 immunofluorescence intensity normalized to the 20-μm diameter intensity for each experiment. \**p* < 0.05, \*\**p* < 0.01, \*\*\**p* < 0.001, \*\*\*\**p* < 0.0001. *N* ≥ 67 images per group for cell area, across 7 independent experiments. *N* ≥ 89 nuclei per group for nuclear area, across three independent experiments. *N* ≥ 73 nuclei per group for micropatterning, based on three independent experiments.

Since cell-cell contacts may also contribute to cell density-dependent LAC-Ig1 immunolabeling in serial dilution seeding experiments, we seeded human fibroblasts as single cells on circular fibronectin micropatterns of either 20 μm or 40 μm in diameter to determine whether cellular and nuclear spreading, independent of cell-cell contacts, could affect LAC-Ig1 immunolabeling. Human fibroblasts adhered to the micropatterns as single cells and spread over the available micropatterned area (Fig. 4C), resulting in cells with either 20- or 40-μm diameter (corresponding to cell spread areas of ≈314 µm^2^and ≈1257 µm^2^, respectively. Nuclear cross-sectional areas were significantly increased on the 40-μm diameter micropatterns compared to the 20-μm diameter micropatterns (Fig. 4D), confirming that increased cell spreading leads to increased nuclear cross-sectional area. Similar to what we had observed in the serial dilution experiments, cells seeded on 40-μm diameter micropatterns had significantly increased LAC-Ig1 immunolabeling compared to cells seeded on 20-μm diameter micropatterns (Fig. 4E), suggesting that differential cell spreading is sufficient to induce Ig-fold conformational changes that results in differential LAC-Ig1 immunolabeling, independent of cell-cell contacts.

### Density dependent Lamin A/C epitope recognition is independent of the apico-basal polarization of the lamina

In another instance of cell context-dependent epitope binding of Lamin A/C antibodies, Vogel and colleagues had found that an antibody directed against an epitope within the Lamin A/C Ig-fold produced labeling with apico-basal polarization that was more pronounced when cells were able to spread or grown on rigid substrates (Ihalainen et al., 2015). Since we observed similar cell spreading-dependent effects in the LAC-Ig1 labeling, we set out to determine if the two phenomena shared a common mechanism, and whether apico-basal polarization was also dependent on cell density. We examined apico-basal polarization of Lamin A/C in human fibroblasts seeded at three representative densities (0.0625×, 0.25×, 1.0×). Consistent with the previous report (Ihalainen et al., 2015), we found that immunofluorescence labeling with the LAC-Ig1 antibody produced a characteristic apico-basal distribution, with the apical nuclear lamina brightly labeled, whereas the basal side hardly showed any fluorescence (**Fig. S6A**). In contrast to Lamin A/C, Lamin B1 showed little apico-basal polarization, as previously reported (Ihalainen et al., 2015). To quantify the apico-basal polarization, we divided each nucleus into the vertical top and bottom halves, summing the fluorescence intensity across each half, and taking the ratio of basal:apical fluorescence intensity. Human fibroblasts showed a similarly polarized ratio of basal:apical LAC-Ig1 fluorescence intensity regardless, of cell density (**Fig. S6B**). The Lamin B1 basal:apical fluorescence intensity ratio was similar across all seeding densities (**Fig. S6C**). Taken together, these results indicate that although we also observed cell density dependent differences in cell spreading and nuclear flattening, the cell density-dependent LAC-Ig1 labeling was independent of apico-basal difference in Lamin A/C organization and must result from a different mechanism.

Lamin A/C are found both at the nuclear periphery as part of the nuclear lamina, and within the nucleoplasm, reflecting differences in mobility and binding partner interaction (Zwerger et al., 2015; Takeshi et al., 2016; Naetar et al., 2021). To address whether the cell density dependent epitope recognition could be sensitive to differences between nucleoplasmic and peripheral Lamin A/C, we quantified the intranuclear distribution of LAC-Ig1 immunofluorescence labeling, comparing nucleoplasmic and nuclear periphery signals in cells seeded at either high (1.0×) or low (0.0625×) cell density (**Fig. S7A**). The high resolution images of confocal cross-sections (**Fig. S7A**) did not indicate obvious differences in the intranuclear distribution of Lamin A/C as a function of high or low cell density. Cells at low density had significantly increased LAC-Ig1 immunofluorescence intensity both in the nucleoplasm and at the nuclear periphery (**Fig. S7B**), consistent with our above findings of the overall immunofluorescence labeling. Differences in the nucleoplasmic to peripheral LAC-Ig1 labeling were not quite statistically significant (*p* = 0.06) between the high and low cell density and overall lower than those of the individual immunofluorescence intensity values (**Fig. S7C**). Since the immunofluorescence intensities for the high cell density condition, particularly for nucleoplasmic Lamin A/C, were often quite low, the ratio of nucleoplasmic to peripheral lamins at these conditions may thus be more prone to variation, and the ratiometric results should be interpreted with some caution.

### Density-dependent epitope binding is not regulated by actin or microtubule polymerization

Previous studies reported that mechanoresponsive conformational changes in the Lamin A/C-Ig fold were mediated by the actin cytoskeleton, particularly through apico-basal compression of the nucleus by apical actin stress fibers (Swift et al., 2013; Ihalainen et al., 2015). To determine whether cell density-dependent Lamin A/C Ig-fold epitope recognition could similarly be related to differing actin filament (F-actin) levels and organization at different seeding densities, we quantified levels of F-actin and LAC-Ig1 binding at different cell densities, and examined whether depolymerization of F-actin using Cytochalasin D abolished density-dependent differences in LAC-Ig1 labeling. Although cells at lower cell densities had (as expected) significantly more F-actin per cell (Fig. 5A, B), disruption of F-actin did not eliminate the difference in LAC-Ig1 immunofluorescence labeling between low and high cell densities (Fig. 5C, D). Levels of Lamin B1 did not change with cell density or cytoskeletal disruption (Fig. 5E). These findings suggest that that forces exerted on the nucleus from the actin cytoskeleton are insufficient to explain density-dependent LAC-Ig1 labeling.

**Figure 5.**
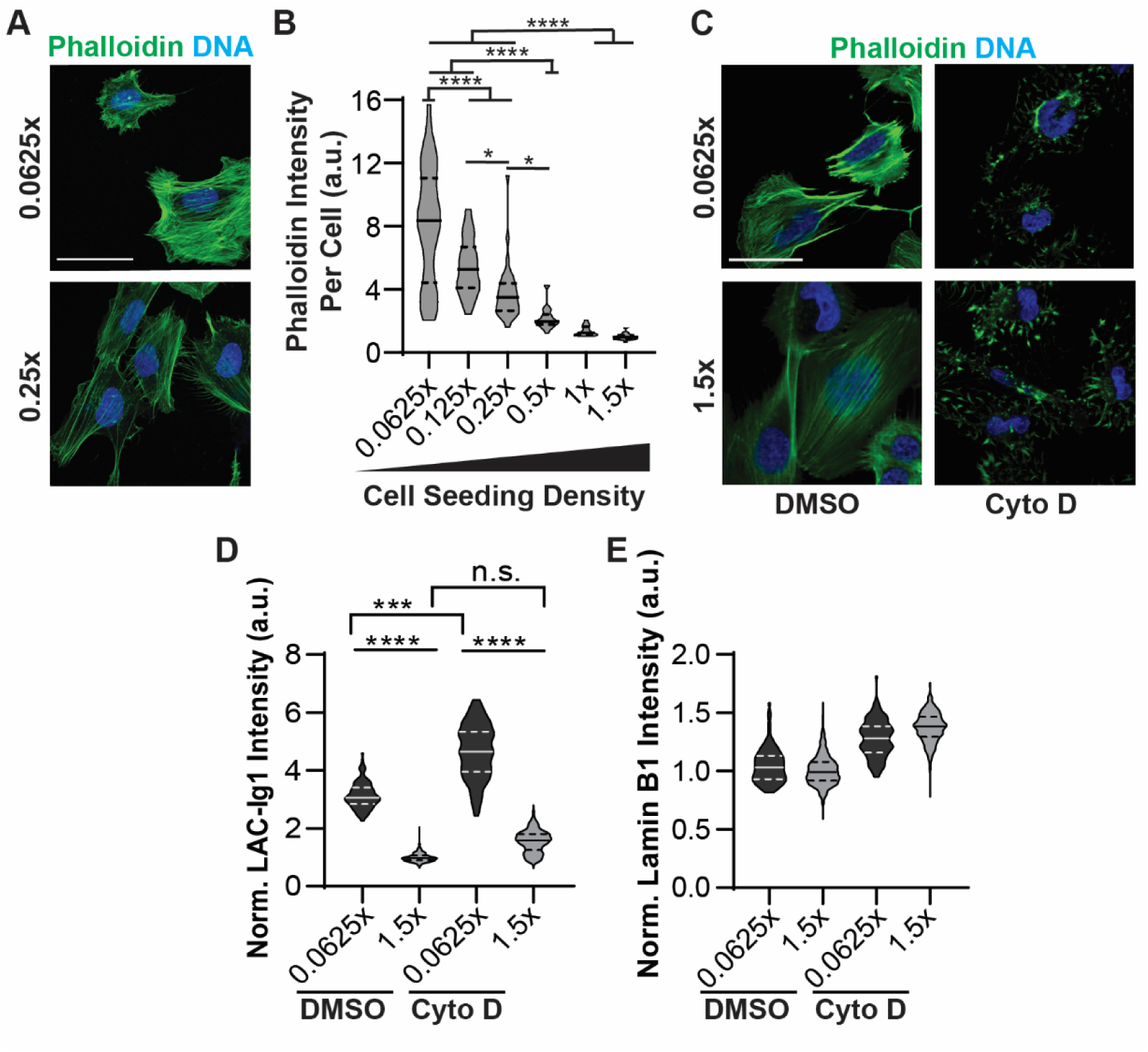
Increased F-actin at low cell densities does not explain cell density dependent binding of LAC-Ig1 antibody. (**A**) Representative image of human fibroblasts stained for F-actin (Phalloidin) and DNA (DAPI). Scale bar: 50 µ m. (**B**) Quantification of the average Phalloidin fluorescence intensity per cell at different densities, showing a significant increase at low cell densities. *, *p* < 0.05; ****, *p* < 0.0001. n.s., non-significant; *N* ≥ 30 images per group, across three independent experiments. (**C**) Representative images of human fibroblasts treated with Cytochalasin D (Cyto D) or vehicle control (DMSO), showing substantial disruption of F-actin in the Cytochalasin D treated cells. Scale bar: 50 µm. (**D**) Quantification of LAC-Ig1 immunofluorescence intensity at low (0.0625×) and high (1.5×) cell densities, in the absence or presence of Cytochalasin D treatment. ****, *p* < 0.0001. *N* ≥ 30 cells per group, across three independent experiments. (E) Quantification of lamin B1 immunofluorescence at low (0.0625×) and high (1.5×) cell densities, in the absence or presence of Cytochalasin D treatment. *N* ≥ 100 cells per group, across three independent experiments. None of the differences were statistically significant.

While the LAC-Ig1 density-dependent immunolabeling was not ablated by disrupting F-actin, other cellular components may also exert mechanical forces on the nuclear envelope and thereby induce changes of the Ig-fold. To determine whether microtubules or microtubule associate motors mediate cell density dependent Lamin A/C Ig-fold recognition, we depolymerized microtubules using Nocodazole and quantified LAC-Ig1 immunofluorescence intensity (**Fig. S8A**). Neither microtubule disruption nor combined treatment with Nocodazole and Cytochalasin D eliminated the density-dependent labeling effect of LAC-Ig1 (**Fig. S8A**). In an alternative approach, we tested whether disrupting the LINC complex, which is crucial to transmit forces between the nucleus and cytoskeleton (Crisp et al., 2006; Lombardi et al., 2011), could eliminate the cell density dependent LAC-Ig1 labeling effect. However, LINC complex disruption using a dominant-negative KASH domain construct (DN-KASH) did not abolish the difference in LAC-Ig1 immunolabeling in human fibroblasts seeded at the low versus high cell densities, nor did LINC complex disruption change the LAC-Ig1 fluorescence intensity compared to mock controls (**Fig. S8B**). Taken together, these findings suggest that the density-dependent LAC-Ig1 labeling is independent of cytoskeletal forces on the nucleus.

Since chromatin plays an important role in governing nuclear mechanics (Stephens et al., 2017, 2018b; Stephens, 2020), chromatin compaction may mediate Lamin A/C Ig-fold conformational changes or interaction with binding partners. Thus, we examined whether inhibiting histone deacetylases with Trichostatin A (TSA) to decrease chromatin compaction could alter cell density dependent LAC-Ig1 binding. Although TSA treatment increased LAC- Ig1 immunolabeling compared to vehicle controls, it did not eliminate the significant difference in LAC-Ig1 fluorescence intensity between low and high cell seeding densities (**Fig. S8C**).

### Density-dependent epitope binding effect is restricted to a narrow region in the Lamin A/C Ig-fold

To gain further insights into the cell density-dependent antibody binding, we performed additional experiments with human fibroblasts seeded at different cell densities and immunofluorescently labeled with three different antibodies against Lamin A/C: two antibodies (LAC-Ig2 and LAC-Ig3) recognizing regions of the Ig-fold that partially overlap with the LAC- Ig1 antibody but have more precisely identified epitopes (Dyer et al., 1997; Ihalainen et al., 2015), and another antibody (LAC-N) binding the Lamin A/C N-terminus (Fig. 6A). The Lamin A/C-Ig2 antibody had previously been shown by Vogel and colleagues to exhibit a differential apico-basal accessibility that responds to changes in nuclear tension and compression (Ihalainen et al., 2015). Although showing a slight trend towards reduced labeling at the highest cell density, the LAC-N antibody did not exhibited any significant changes in immunolabeling across different seeding densities (Fig. 6B), suggesting that cell density primarily affects antibody recognition of the Ig-fold epitope. Interestingly, the LAC-Ig2 antibody showed only a weak cell density dependent effect in its labeling, with only the lowest cell density reaching a statistically significant difference compared to the highest cell density (Fig. 6C). This finding suggest that the epitope recognized by the LAC-Ig2 antibody is less affected by cell density than the epitope targeted by the LAC-Ig1 antibody. However, we found that the LAC-Ig2 labeling was extremely variable and appeared very sensitive to the fixation conditions. The immunofluorescence intensity varied strongly between experiments, with some producing labeling that was hardly detectable, and others showing a strong inverse correlation between the cell density and the LAC-Ig2 immunofluorescence intensity (**Fig. S9**). Since the LAC-Ig2 antibody is sensitive to Lamin A/C conformational changes (Ihalainen et al., 2015), we hypothesize that slight variations in the fixation conditions may arrest the corresponding epitope in different conformations or affect interaction with Lamin A/C binding partners that mask the epitope. This experiment-to-experiment variability may potentially conceal a stronger effect of density-dependent immunolabeling with the LAC-Ig2 antibody. Surprisingly, the LAC-Ig3 antibody, which binds a specific region of the Lamin A/C Ig-fold containing amino acids 476-484, although residues 477- 478 dominate (Manilal et al., 2004), thus overlapping with both the LAC-Ig1 and LAC-Ig2 antibodies, did not show any cell density-dependent changes in immunofluorescence labeling (Fig. 6D), suggesting that these specific region of the Lamin A/C Ig-fold is not affected by density-dependent epitope changes.

**Figure 6.**
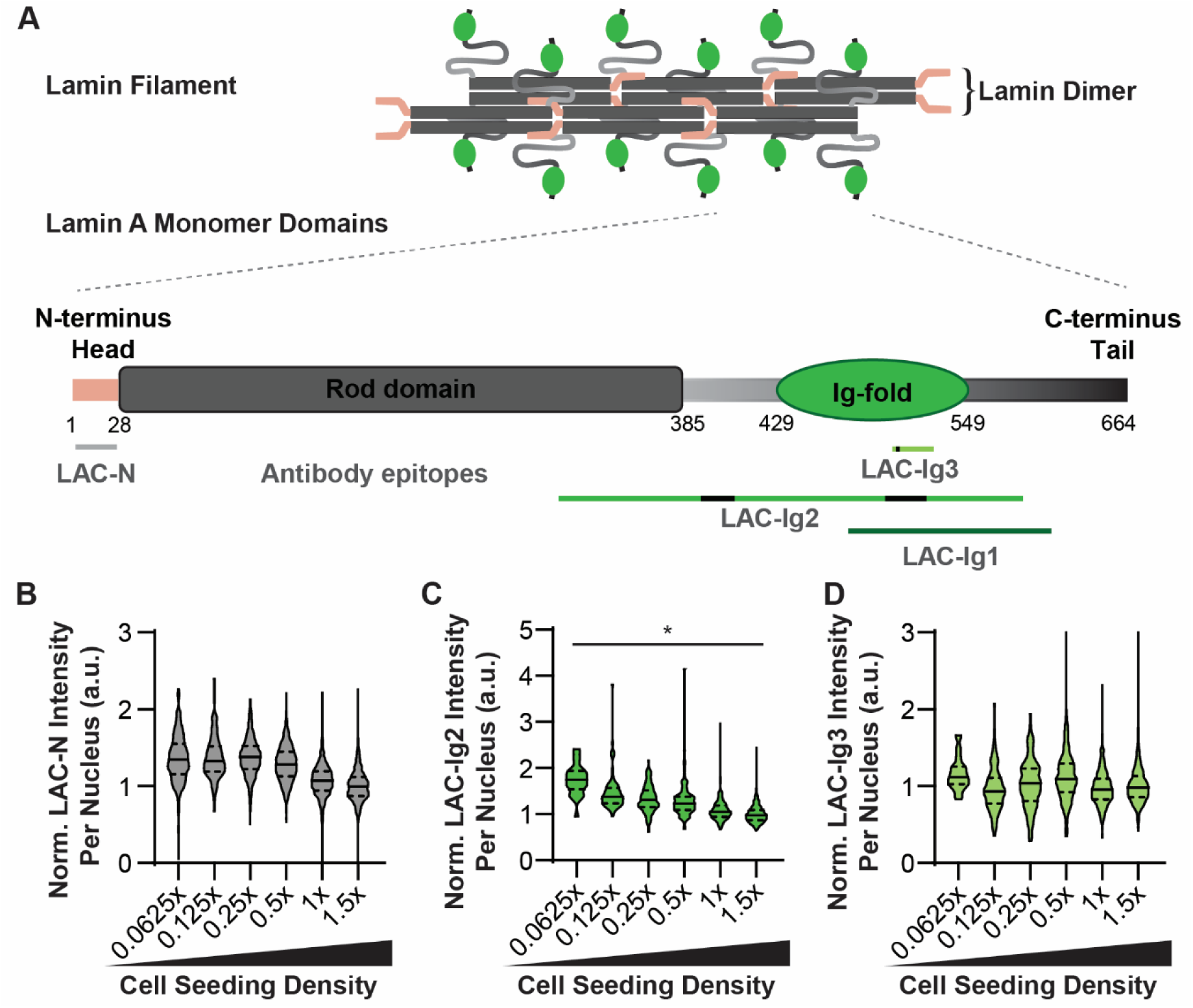
Density-dependent antibody binding is highly epitope specific. (**A**) Schematic overview of Lamin A protein structure, both as monomer and assembled into filaments (Turgay and Medalia, 2017; Turgay et al., 2017; Ahn et al., 2019), indicating the specific binding sites of the four Lamin A/C antibodies used in this study: the LAC-Ig1 antibody, two additional antibodies targeting the Lamin A/C Ig-fold (LAC-Ig2 and LAC-Ig3) whose epitopes (partially) overlap the postulated binding region of the LAC-Ig1 antibody, and an antibody binding the N-terminus of Lamin A/C (LAC-N). The antibodies have been shown to bind generally the regions marked with green lines, with the strongest/specific sites from epitope mapping shown as black segments. This specific epitope mapping has not been done for the LAC- Ig1 antibody. (**B**) Quantification of LAC-N immunofluorescence labeling at different cell densities. The LAC-N antibody did not show any cell density-dependent differences in labeling. (**C**) Quantification of LAC-Ig2 immunofluorescence labeling at different cell densities. The LAC-Ig2 antibody shows a weak inverse trend between cell seeding density and normalized fluorescence intensity. \**p* < 0.05. *N* > 67 nuclei per group, based on three independent experiments. (**D**) Quantification of LAC-Ig3 immunofluorescence labeling at different cell densities. The Lamin A/C-Ig3 antibody did not exhibit any cell density-dependent differences in fluorescence intensity.

Taken together, the density-dependent epitope binding of the LAC-Ig1 and LAC-Ig2 antibodies suggest that the Ig-fold region encompassing these epitopes is particularly sensitive to conformational or other changes associated with different cell densities, such as interaction with Lamin A/C binding partners that mask the epitope (our present work) or cytoskeletal forces within the cell (Ihalainen et al., 2015), thus explaining the conditional epitope binding. The LAC-Ig2 antibody has been precisely epitope-mapped, showing that it binds Lamin A/C at amino acids 475-497 (Ihalainen et al., 2015), which falls in the C’E loop of the Ig-fold (Fig. 7A) (Krimm et al., 2002). The cryptic Cys^522^ residue, which can become exposed in response to mechanical stress (Swift et al., 2013), falls in another loop domain of the Ig-fold, the EF loop (Fig. 7A). Although the specific binding sites of the LAC-Ig1 antibody has not been mapped yet, the general region it is known to bind overlaps the surface of the Ig-fold (Manilal et al., 2004), including both the C’E and EF loop regions (Fig. 7A-B). Therefore, we hypothesize that the LAC-Ig1 antibody binds the C’E and/or the EF loop of the Ig-fold, which undergoes conformational changes in response to different cell densities through a yet to be determined mechanism, thereby modulating epitope recognition by the LAC-Ig1 antibody, and to lesser degree the LAC-Ig2 antibody. Another, non-mutually exclusive potential mechanism is that the epitope may become masked by cell density dependent interaction of the Lamin A/C Ig-fold with one or more binding partner(s) that increases with higher density. As the Ig-fold interacts with numerous binding partners, including emerin, MAN1, nuclear pore proteins, and DNA (Ho and Lammerding, 2012; Donnaloja et al., 2020) whose interactions may similarly affected by the cell density dependent conformational changes or epitope masking, this opens the intriguing possibility of the Lamin A/C Ig-fold serving as cell density sensor.

**Figure 7.**
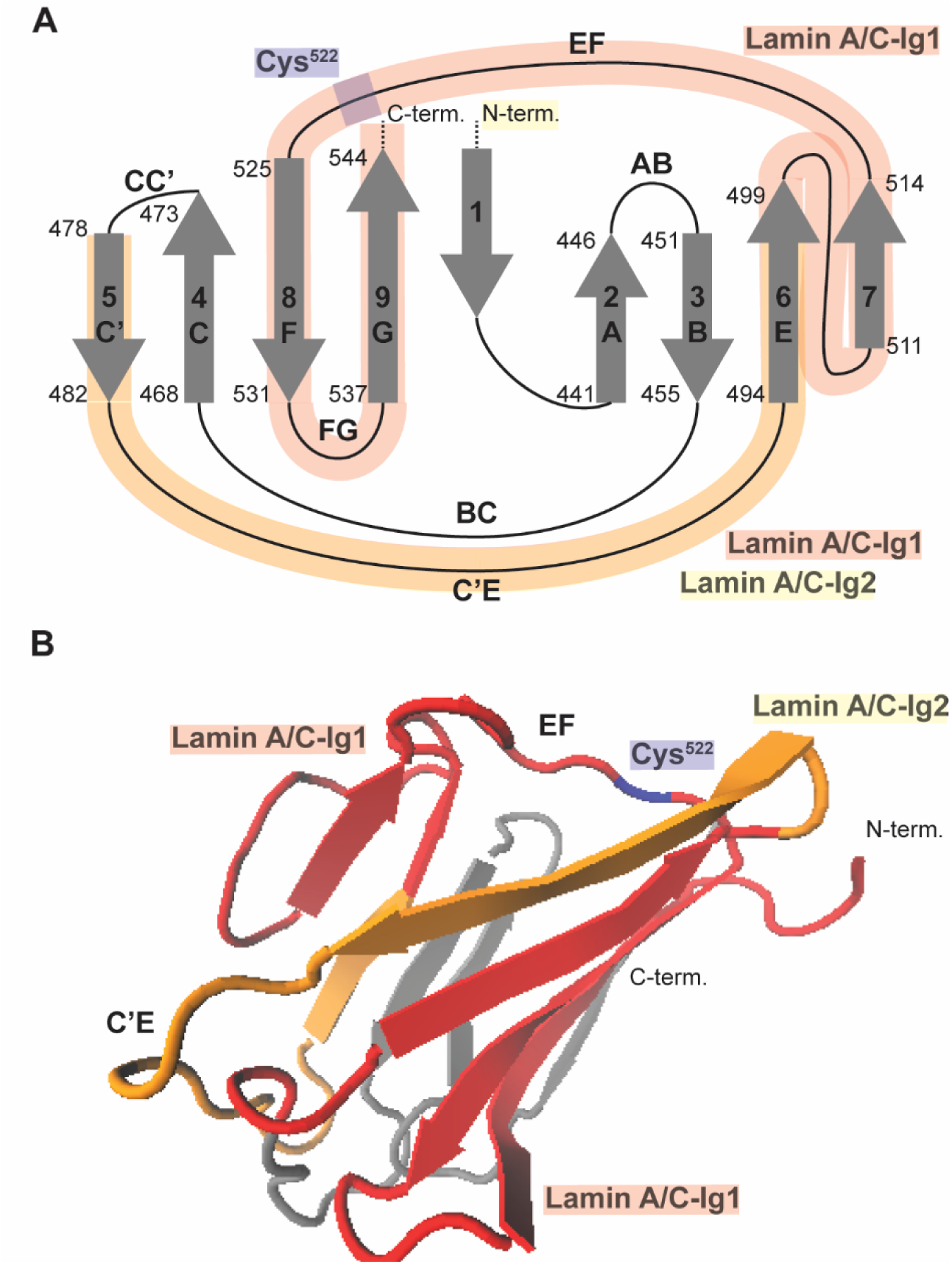
Lamin A/C-Ig1 antibody binding in the Ig-fold structure. (**A**) Schematic map of the Lamin A/C Ig-fold protein structure. Regions known to be mechanosensitive, based on the reported differential LAC-Ig2 epitope binding (Ihalainen, *et al*, 2015) and exposure of the cryptic Cys^522^ residue in response to shear stress (Swift, *et al*, 2013), are highlighted in yellow and blue, respectively. The regions recognized by the Lamin A/C-Ig1 antibody are shaded in red, notably overlapping two loop regions – C’E and EF – that are known to be mechanosensitive. (**B**) Ribbon diagram of the Lamin A/C Ig-fold structure, in which the postulated regions recognized by the LAC-Ig1 antibody are shaded in red, notably overlapping the two mechanoresponsive loop regions, C’E and EF.

## Discussion

The Lamin A/C Ig-fold has been shown previously to undergo mechanically induced conformational changes in response to cellular shear stress or local force transmission from apical actin fibers to the nuclear envelope (Swift et al., 2013; Ihalainen et al., 2015), which can result in variable immunofluorescence labeling by antibody targeting epitopes within the Ig-fold (Ihalainen et al., 2015). Here, we demonstrated that differential labeling with Ig-fold targeting antibodies can also result from changes in cell density. This effect was particularly pronounced for a commonly used antibody (JOL-2; referred to here as LAC-Ig1), which had not been recognized previously to be sensitive to mechanically induced changes. The immunofluorescence intensity was about 4-times higher at low cell densities compared to high cell densities (Fig. 1D), even when actual Lamin A/C levels did not change, as assessed by Western analysis or fluorescently tagging endogenous Lamin A/C. Lamin A/C is a commonly used marker for the nucleus, and the quantification of Lamin A/C levels an important biomarker for countless physiological phenomenon, including development, stem cell differentiation, cancer, and aging, among others (Rober et al., 1989; Constantinescu et al., 2006; Duque and Rivas, 2006; Willis et al., 2008; Irianto et al., 2016; Dubik and Mai, 2020; Bell et al., 2021). Furthermore, changes in Lamin A/C levels are increasingly referred to as indicators of nuclear mechanotransduction and cellular mechano-responsiveness (Swift et al., 2013; Ihalainen et al., 2015; Iyer et al., 2021). As such, the results presented here indicate that it is critical for studies using the LAC-Ig1 (JOL-2), LAC-Ig2 (131C3), and likely other Lamin A/C Ig-fold targeting antibodies that recognize loop regions of the protein structure, to be aware of the sensitivity of these antibodies to cell density.

While our results share some similarities with the previous reports of conformational changes in the Lamin A/C Ig-fold (Swift et al., 2013; Ihalainen et al., 2015), including a sensitivity to cellular spreading, there are several note-worthy distinctions between our findings and those previous studies. First, whereas the mechanoresponsive conformational changes responsible for the differential apico-basal immunofluorescence labeling required apical actin stress fibers and intact LINC complexes (Ihalainen et al., 2015), the density-dependent labeling with the LAC-Ig1 persisted even when depolymerizing the actin cytoskeleton or disrupting the LINC complex. Second, despite some overlap of the epitopes recognized by the LAC-Ig1 and the LAC-Ig2 antibodies, we observed only a modest apico-basal polarization in the binding of the LAC-Ig1 antibody compared to that reported previously for the LAC-Ig2 antibody (Ihalainen et al., 2015). The apico-basal polarization did not vary with cell seeding density, further supporting the idea that the differential apico-basal antibody binding and the density dependent binding result from separate mechanisms. Third, the density dependent effect was limited to a very specific epitope, as another antibody recognizing the Ig-fold, LAC-Ig3, did not exhibit any variations in labeling Lamin A/C at different cell densities (Fig. 6D).

One limitation of the current study is that the mechanism responsible for the density dependent antibody binding remains unclear. Based on the overlap of the affected Ig-fold region with previously recognized conformationally variable domains of the Ig-fold (Swift et al., 2013; Ihalainen et al., 2015), the location of the region at the interface of the C’E and EF loop regions, and the sensitivity of the LAC-Ig2 antibody to fixation conditions, we envision a scenario in which this Ig-fold region is particularly prone to conformational changes that alter the affinity of the antibodies targeting this epitope, thus affecting antibody recognition and labeling. However, which specific stimulus causes such a conformation change in response to increasing cell density remains to be determined. Given that neither actin, microtubule, nor LINC complex disruption abolished the density-dependent effect, it appears unlikely that the effect is directly triggered by mechanical forces on the nucleus. An alternative explanation may thus be that cell density dependent interaction of one or more Lamin A/C binding partners with the identified Ig-fold region results in epitope masking at high cell density, leading to the reduced immunofluorescence labeling. The exact identity of the binding partner(s), and how cell density modulates the interactions, remain to be elucidated. Although an increase in cell-cell contacts could be a potential explanation, our experiments with single cells seeded on micropatterns of different sizes, which resulted in size-dependent LAC-Ig1 immunofluorescence labeling (Fig. 4C-E), indicate that the effect must be at least in part independent of cell-cell contacts.

Another limitation of the study is that the precise epitope recognized by the LAC-Ig1 antibody has not been determined yet. While outside the scope of the current study, a more precise mapping of the epitope could aid in narrowing down the specific Ig-fold region responsible for the density-dependent effect.

Furthermore, we were unable to detect specific functional consequences associated with the altered antibody recognition. Neither nuclear stiffness nor force transmission between the cytoskeleton and nucleus varied with cell density (Fig. 3). Nonetheless, given the diverse functions of Lamin A/C and the numerous proteins that can interact with the Ig-fold, one can easily envision how density-dependent conformational changes or protein-protein interaction of the Lamin A/C Ig-fold could modulate various cellular functions. This idea is further supported by the numerous disease causing *LMNA* mutations found within the Ig-fold (Bertrand et al., 2011; Dutta et al., 2018). The Ig-fold is a hotspot for *LMNA* mutations, many of which are located in the C’E and EF loop regions (Krimm et al., 2002) that we hypothesize to be responsible for the density-dependent LAC-Ig1 epitope binding and other mechanosensitive behaviors (Swift et al., 2013; Ihalainen et al., 2015). *LMNA* mutations occurring in the Ig-fold are known to destabilize the Ig-fold structure (Krimm et al., 2002; Bera et al., 2014; Dutta et al., 2018) and cause changes in nuclear organization (Vigouroux et al., 2001; Bera et al., 2014) and chromatin configuration (Vigouroux et al., 2001; Verstraeten et al., 2009). Such changes are already known to disrupt nuclear mechanotransduction processes and hence downstream biochemical signaling pathways (Davidson and Lammerding, 2014; Maurer and Lammerding, 2019; Donnaloja et al., 2020). *LMNA* mutations may cause changes in the Ig-fold protein structure that could disrupt the normal conformational changes and/or altered interaction with binding partners of the Ig-fold in response to mechanical or biochemical signals, thereby causing improper adaptation of the nucleus to the cell’s microenvironment. As more information becomes available about both the Ig-fold’s adaptation to external stimuli and how *LMNA* mutations, particularly those falling in Ig-fold loop regions, disrupt normal cellular function, this link should be explored further in future studies.

In conclusion, this study demonstrates a previously unrecognized cell-density dependent effect of the interaction of the Lamin A/C Ig-fold with commonly used antibodies that can dramatically alter immunofluorescence measurements, and that indicate intriguing underlying changes in Ig-fold conformation or binding partner interaction involving the Ig-fold loop regions. These density-dependent changes may be critical for regulating interactions with the many Ig-fold binding partners of this region that are involved in various cellular functions and represent a path for further investigation in both *LMNA* wild-type and mutant cells.

## Acknowledgements

We thank the Biotechnology Resource Center (BRC) Flow Cytometry Facility (RRID:SCR_021740) and sequencing facility (RRID:SCR_021727) at the Cornell Institute of Biotechnology for their resources and technical assistance. This work was performed in part at the Cornell NanoScale Science & Technology Facility, a member of the National Nanotechnology Coordinated Infrastructure, which is supported by the National Science Foundation (award NNCI-2025233). This work was supported by awards from the National Institutes of Health (R01 HL082792, R01 GM137605, U54 CA210184 to J.L.), the National Science Foundation (CBET 1715606, MCB-1715606, and URoL-2022048 to J.L.; Graduate Research Fellowships 2016229710 to M.W. and 2014163403 to G.R.F.), the Department of Defense Breast Cancer Research Program (Breakthrough Award BC150580 to J.L.), the American Heart Association (20PRE35080179 to M.M.), and the Volkswagen Foundation (A130142 to J.L.). The content of this manuscript is solely the responsibility of the authors and does not necessarily represent the official views of the National Institutes of Health.

## Data Availability

The authors confirm that the data supporting the findings of this study are available within the article and its supplementary materials.

