## Supplementary Materials for "The Lamin A/C Ig-fold undergoes cell density-dependent changes that alter epitope binding"

### <u>Supplementary Figures</u>

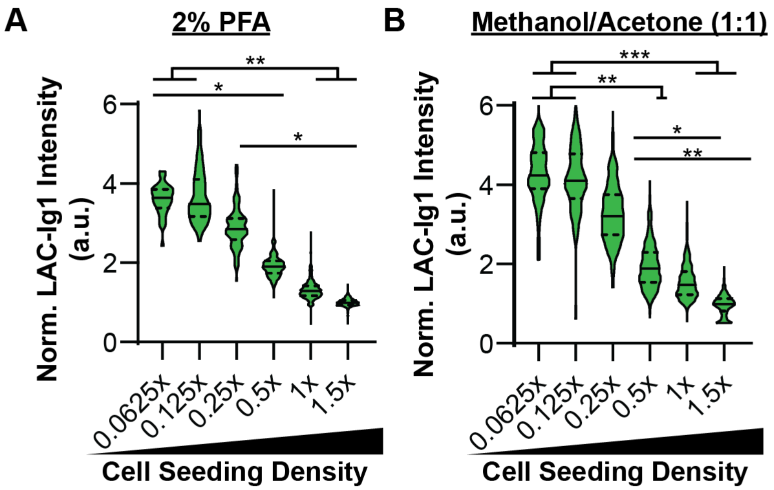


**Figure S1. Lamin A/C-Ig1 density-dependent immunolabeling is independent of fixation method.** Quantification of LAC-Ig1 immunofluorescence intensity of human fibroblasts seeded at different cell densities and fixed with either (**A**) 2% paraformaldehyde (PFA) or (**B**) 1:1 Methanol:Acetone. Both fixation methods produced similar inverse relationships between LAC-Ig1 fluorescence intensity and cell seeding density. *, $p < 0.05$; **, $p < 0.01$; ***, $p < 0.001$. $N >$ 90 nuclei per group from three independent experiments.

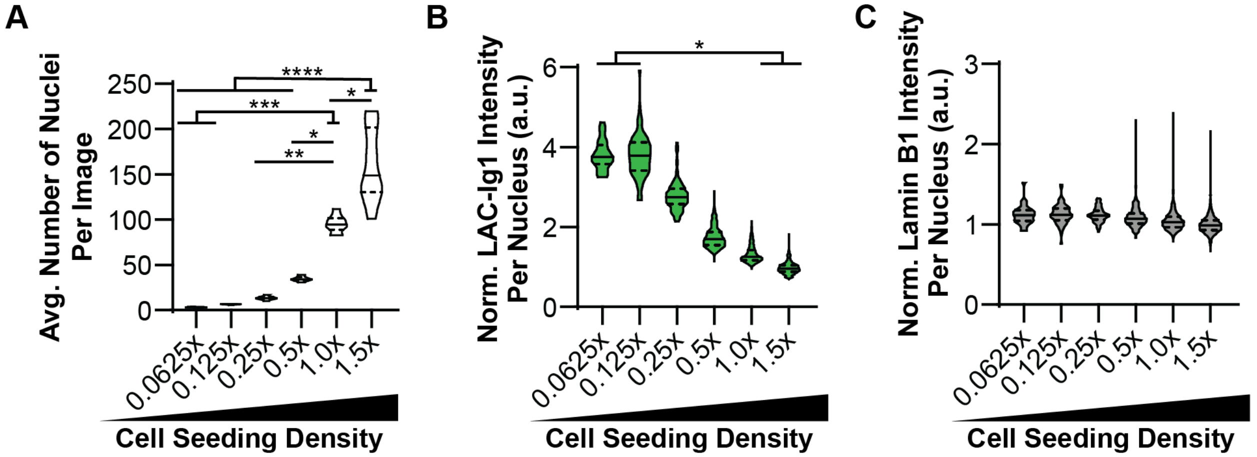


**Figure S2. HeLa cells show an inverse relationship between Lamin A/C-Ig1 fluorescence intensity and cell seeding density.** (**A**) Average number of HeLa nuclei per image shows an exponential trend, as expected. *, $p < 0.05$; **$p < 0.01$; ***, $p < 0.001$; ****, $p < 0.0001$. $N = 30$ images per group for number of nuclei, collected from three independent experiments. (**B**) LAC-Ig1 immunofluorescence intensity normalized to the levels at the highest cell seeding density, 1.5×, varies inversely with cell seeding density. *, $p < 0.05$. $N > 100$ nuclei per group, pooled from three independent experiments. (**C**) Lamin B1 immunofluorescence intensity normalized to the highest cell seeding density, 1.5×. Differences between groups were not statistically significant. $N > 100$ nuclei per group, based on three independent experiments.

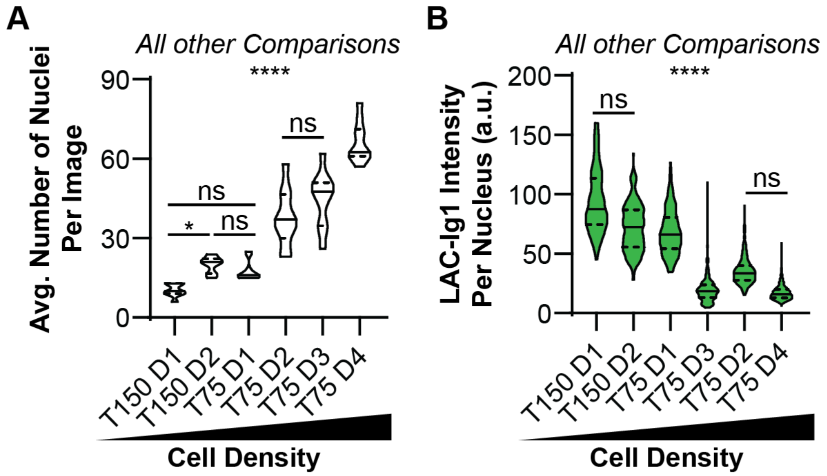


**Figure S3. MDA-MB-231 cells seeded at similar density but grown to different densities exhibit density-dependent LAC-Ig1 immunolabeling.** (**A**) Quantification of cell density after growth for one to four days (D1 – D4) for cells initially seeded at similar cell density. The average number of MDA-MB-231 nuclei per image upon growth over several days followed a similar trend as the serial dilution approach used in other experiments. *, $p < 0.05$; ****, $p < 0.0001$; ns, not statistically significant. Unless otherwise indicated, $p < 0.0001$ for all comparisons. $N = 10$ images per group. (**B**) MDA-MB-231 cells grown over several days show the same significant inverse relationship between Lamin A/C (LAC)-Ig1 fluorescence intensity and cell seeding density. *, $p < 0.05$, ****, $p < 0.0001$. Unless otherwise indicated, $p < 0.0001$ for all comparisons. $N = 10$ images for number of nuclei counts, $N > 100$ nuclei per group.

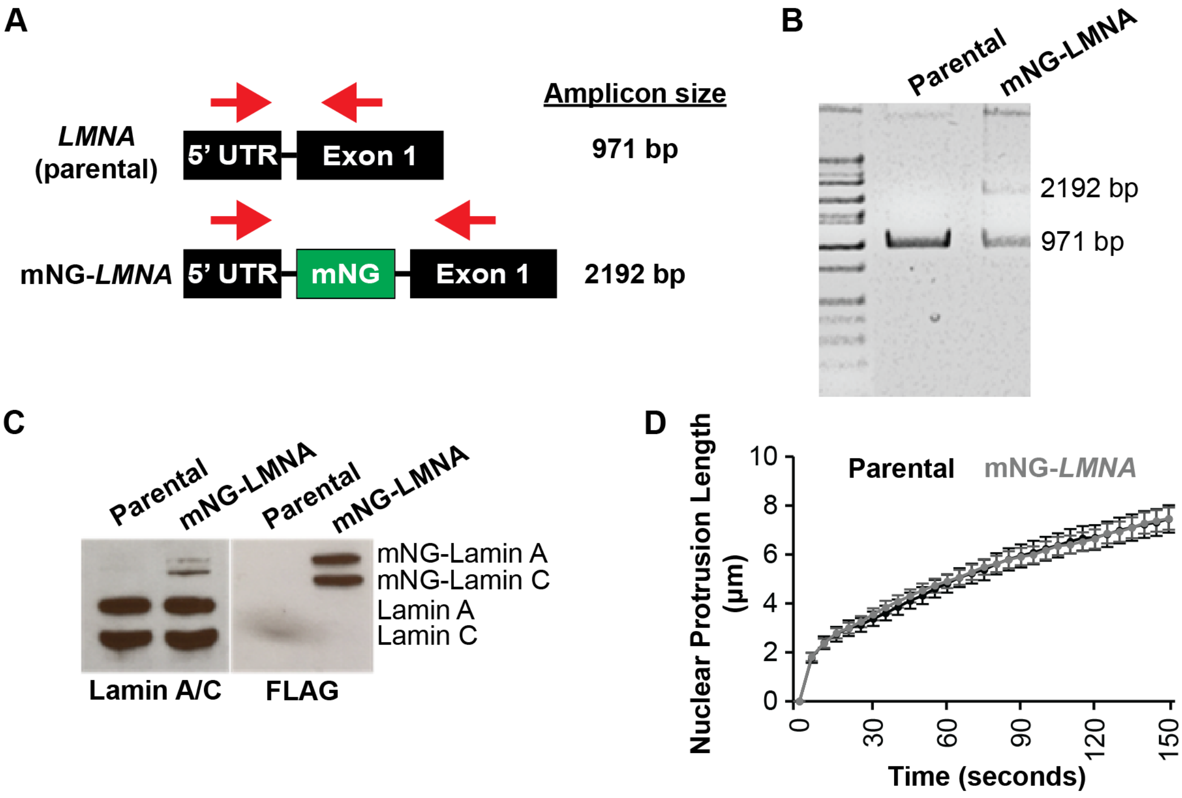


**Figure S4. Design and validation of labeling endogenous lamin A/C with mNeonGreen.** (**A**) Schematic overview of the strategy to insert mNeonGreen (mNG) into the endogenous *LMNA* gene. The sequence encoding mNG was inserted into the *LMNA* gene in between the 5' Untranslated Region (UTR) and Exon 1 of MDA-MB-231 cells, resulting in the expression of an N-terminal mNG-lamin A/C fusion product (mNG-*LMNA*). Red arrows indicate the location and direction of primers designed to confirm that the mNG sequence was successfully inserted. Insertion of the mNG sequence should increase the amplicons size from 971 to 2192 bp. (**B**) PCR products using the primers shown in (A) confirm the presence of a 2192 bp amplicon in mNG-*LMNA* expressing cells. (**C**) Western blot confirming the presence of mNG-Lamin A/C, indicated by the larger size compared to untagged Lamin A/C. The lower expression of the mNG-*LMNA* products suggest that only a single *LMNA* allele was tagged. (**D**) Quantification of nuclear deformability (nuclear protrusion length) of mNG-*LMNA* and untagged parental cells in a microfluidic micropipette aspiration assay. The mNG-*LMNA* cells exhibited similar nuclear mechanics compared to the parental cell line. $N > 48$ nuclei per group for micropipette experiments from two independent experiments.

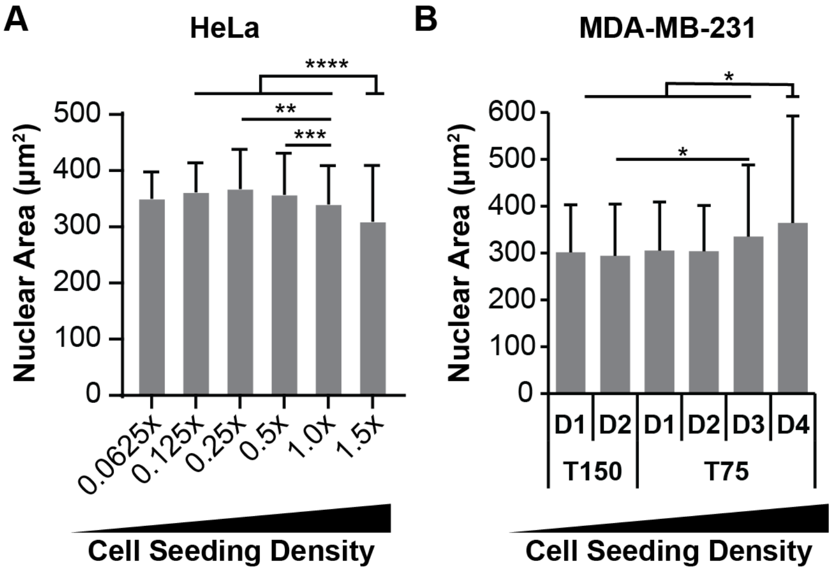


**Figure S5. Nuclear cross-sectional area varies slightly with cell density.** (**A**) Nuclear cross-sectional area of HeLa cells seeded at different cell densities. Cells at the highest density had reduced nuclear cross-sectional area. **, $p < 0.01$; ***, $p < 0.001$; ****, $p < 0.0001$. $N > 50$ nuclei for each group, from three independent experiments. (**B**) Nuclear cross-sectional area of MDA-MB-231 cells seeded at the same low cell density and fixed, stained, and analyzed at different time points to achieve different cell densities at time of fixation). Cells at the highest cell density had increased nuclear cross-sectional area. **, $p < 0.01$. $N > 50$ nuclei for each group, from three independent experiments.

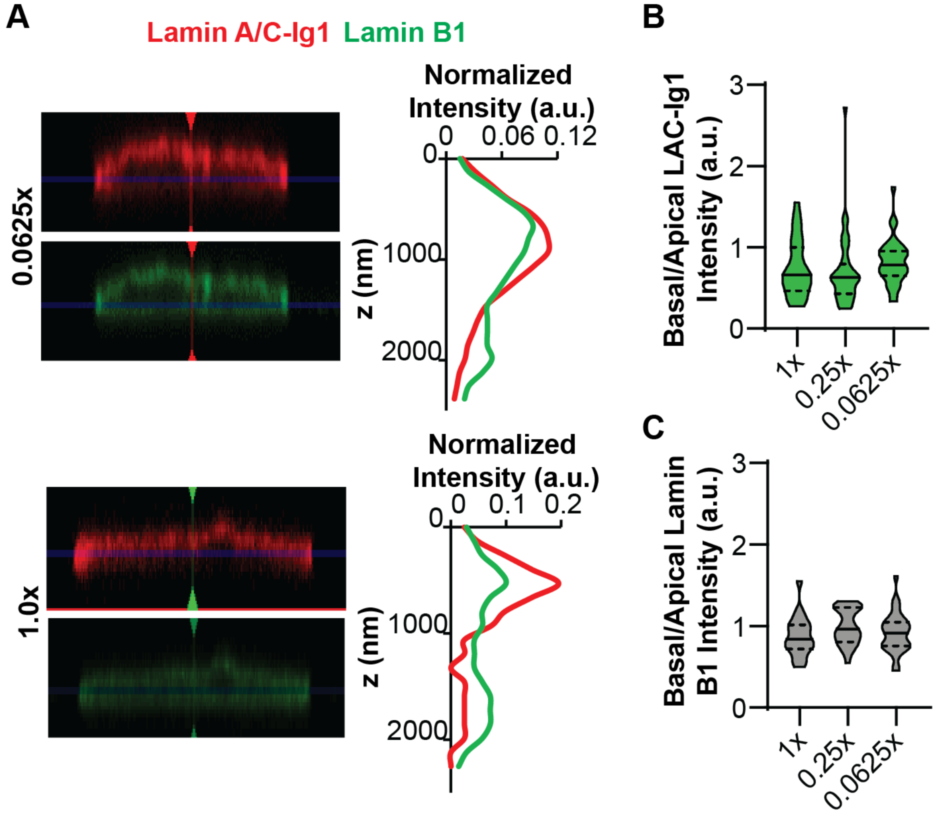


**Figure S6. Apico-basal polarization of LAC-Ig1 labeling is independent of cell density.** (**A**) Representative X-Z cross-sections of human fibroblasts seeded at low (0.0625×) or high (1.0×) cell seeding density that were immunofluorescently labeled with the LAC-Ig1 antibody or an antibody against Lamin B1. Graphs on the right show the corresponding fluorescence intensity profiles in the z-direction, confirming the apico-basal polarization of LAC-Ig1 labeling. (**B**) Quantification of the basal to apical LAC-Ig1 intensity ratio, showing that apico-basal polarization (i.e., a ratio below 1) was present at all seeding densities. (**C**) Quantification of the basal-to-apical Lamin B1 ratio shows lack of apico-basal Lamin B1 polarization in cells at all seeding densities. $N > 20$ nuclei per group from three independent experiments.

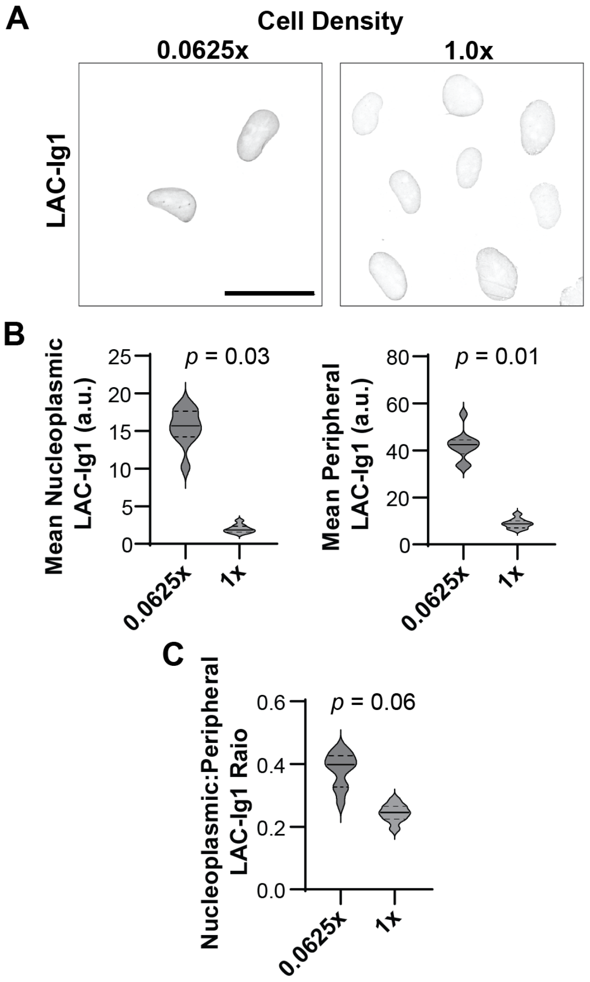


**Figure S7. Intranuclear distribution of Lamin A/C immunofluorescence labeling as a function of cell density.** (**A**) Representative confocal cross-sections of cells seeded either at low (0.0625×) or high (1.0×) cell densities immunofluorescently labeled with the LAC-Ig1 antibody. Scalebar: 50 μm. (**B**) Quantification of the mean nucleoplasmic (left) and peripheral (right) fluorescence intensities for cells at low (0.0625x) and high (1x) cell seeding densities. Both nucleoplasmic and peripheral Lamin A/C staining was significantly higher at low cell density. (**C**) The difference in the ratio of the nucleoplasmic to peripheral LAC-Ig1 immunofluorescence labeling signal was not quite statistically significant between the cell seeding densities. $N \geq 25$ nuclei per group, across three independent experiments.

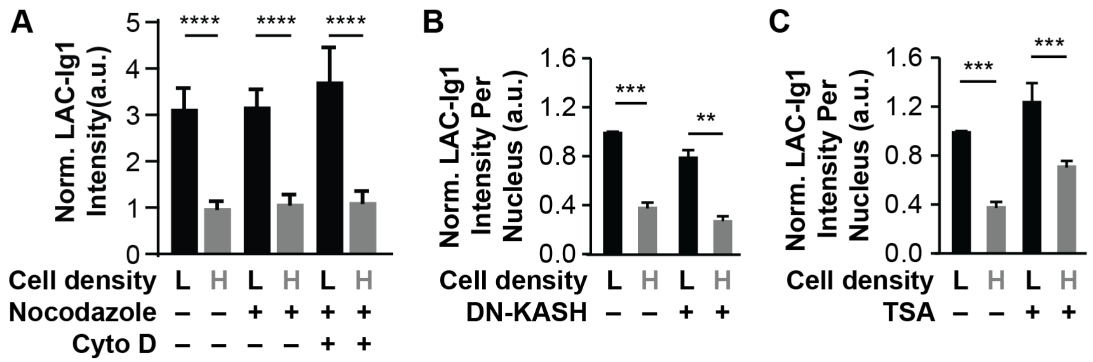


**Figure S8. Density-dependent LAC-Ig1 epitope labeling is independent of cytoskeletal forces or chromatin organization.** (**A**) Quantification of LAC-Ig1 immunofluorescence intensity per nucleus normalized to the low density vehicle control of human fibroblasts seeded at either low (3,125 cells/cm$^2$) or high (31,250 cells/cm$^2$) cell density. Neither treatment with Nocodazole to disrupt microtubules, nor combined treatment with Nocodazole and Cytochalasin D to disrupt microtubules and actin filaments, eliminated the difference in LAC-Ig1 labeling between low and high cell densities. ****, $p < 0.0001$, based on $N > 40$ nuclei per condition, across three independent experiments. (**B**) Disruption of nucleo-cytoskeletal coupling using an inducible dominant-negative KASH construct (DN-KASH) did not change the difference in LAC-Ig1 immunofluorescence intensities between low and the high cell density. **, $p < 0.01$; ***, $p < 0.001$. $N > 45$ nuclei per group for all experiments. (**C**) Treatment with the deacetylase inhibitor Trichostatin A (TSA) increased LAC-Ig1 fluorescence intensity in both high and low seeding densities compared to vehicle controls, but did not eliminate the difference in immunofluorescence labeling between the low and high cell density. ***, $p < 0.001$. $N > 40$ nuclei per group for all experiments, collected in three independent experiments.

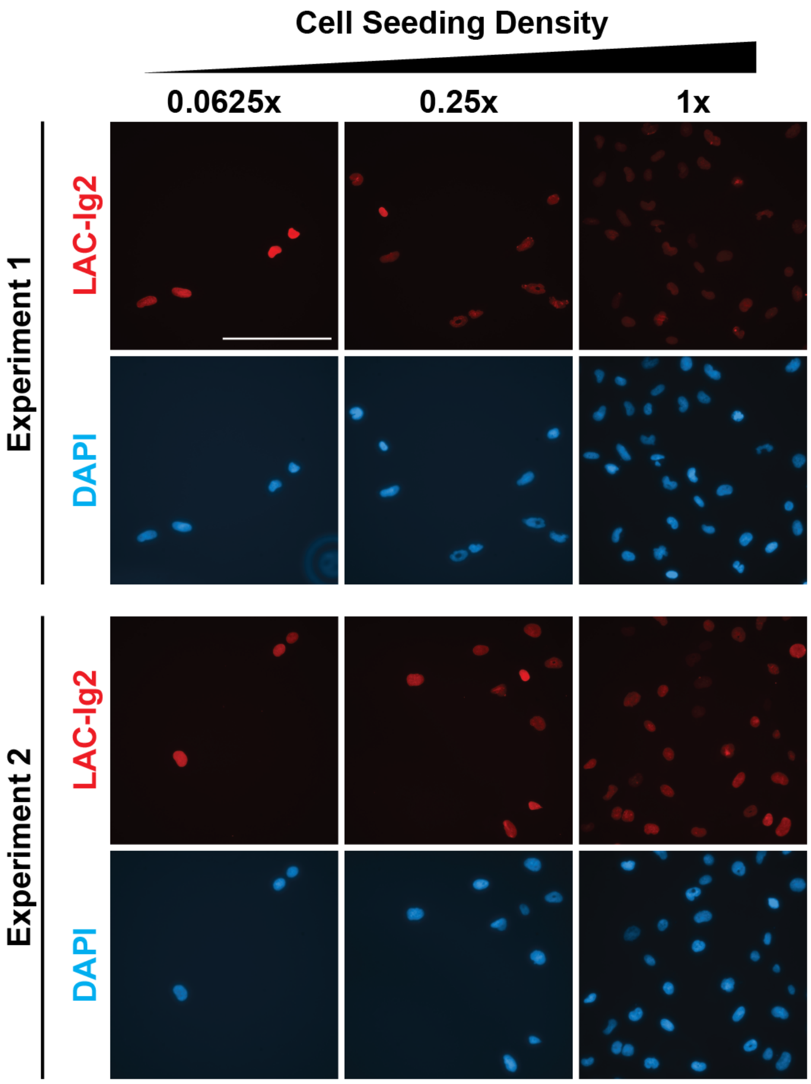


**Figure S9. The LAC-Ig2 antibody has high experiment-to-experiment variability.** Representative immunofluorescence labeling images for LAC-Ig2, along with DAPI staining for DNA, show high variability between experiments in the LAC-Ig2 labeling. Scale bar: 200 µm for all images.

### Supplementary Table(s)

| Antibody | Catalog # | Vendor | Dilution (IF) | Dilution (Western) |
|---|---|---|---|---|
| Lamin A/C JOL-2 (“Ig1”) | ab40567 | Santa Cruz Biotechnologies | 1:200 | 1:2000 |
| Lamin A/C E-1 (“N”) | sc-376248 | Santa Cruz Biotechnologies | 1:150 | N/A |
| Lamin A/C 131C3 (“Ig2”) | ab8984 | Abcam | 1:200 | N/A |
| Lamin A/C 4C10 (“Ig3”) | MANLAC3(4C10) | Developmental Studies Hybridoma Bank | 1:150 | N/A |
| Lamin A/C N-18 | sc-6215 | Santa Cruz Biotechnologies | 1:200 | |
| Lamin B1 | 12987-1-AP | Proteintech | 1:200 | N/A |
| Alpha-tubulin | T9026 | Sigma | | 1:4000 |
| Alexa Fluor 568; Goat anti-mouse IgG1 | A-21124 | Invitrogen | 1:250 | N/A |
| Alexa Fluor 568; Goat anti-mouse IgG2a | A-21134 | Invitrogen | 1:250 | N/A |
| Alexa Fluor 488; Donkey anti-rabbit IgG | A-21206 | Invitrogen | 1:250 | N/A |
| HRP-conjugated antibodies | | Biorad | | 1:1000 |

**Supplementary Table 1.** Primary and secondary antibodies used for immunofluorescence labeling and western analysis.
